# Prevention and reversal of hypertension-induced coronary microvascular dysfunction by a plant-based diet

**DOI:** 10.1101/2025.04.19.649660

**Authors:** Rami S. Najjar, Nedumangalam Hekmatyar, Yanling Wang, Vu Ngo, Hannah L. Lail, Juan P. Tejada, Jessica P. Danh, Desiree Wanders, Rafaela G. Feresin, Puja K. Mehta, Andrew T. Gewirtz

## Abstract

**Background and aims:** Coronary microvascular dysfunction (CMD) is associated with adverse cardiovascular outcomes. CMD is driven by endothelial and vascular smooth muscle cell (VSMC) dysfunction. We aimed to test whether CMD could be mitigated by a plant-based diet (PBD) in an animal model of hypertension.

**Methods:** We compared 28- and 40-week-old female normotensive Wistar-Kyoto and spontaneously hypertensive (SHR) rats, maintained, from age 4 weeks, on a control refined diet or a PBD, comprised of 28% fruits, vegetables, nuts and legumes. A subset of control SHRs were switched to the PBD at 28 weeks. CMD was assessed by coronary flow reserve via echocardiogram. Cardiac microvascular endothelial function was assessed via cMRI. Endothelial and VSMC function were assessed in the left ventricle (LV) or in isolated VSMCs. The role of gut microbiota was probed via 16S sequencing and antibiotics. Cardiac inflammation, oxidative stress, and fibrosis were also explored.

**Results:** SHRs exhibited endothelial dysfunction and likely VSMC dysfunction. PBD did not ameliorate their hypertension but, nonetheless, prevented and reversed CMD. PBD’s mitigation of CMD was associated with improved endothelial nitric oxide synthase function and NO-mediated VSMC signaling, as well as reductions in LV oxidative stress, inflammatory signaling, and fibrosis. PBD altered the gut microbiota, although antibiotic studies failed to establish its importance in ameliorating CMD.

**Conclusions:** A PBD prevented CMD development and reversed established CMD in SHRs. Such benefits of PBD, which occurred without alleviating hypertension, were possibly due to improved endothelial function and likely improved VSMC function. These results support clinical trials to test PBDs in human CMD.

**Graphical Abstract:** 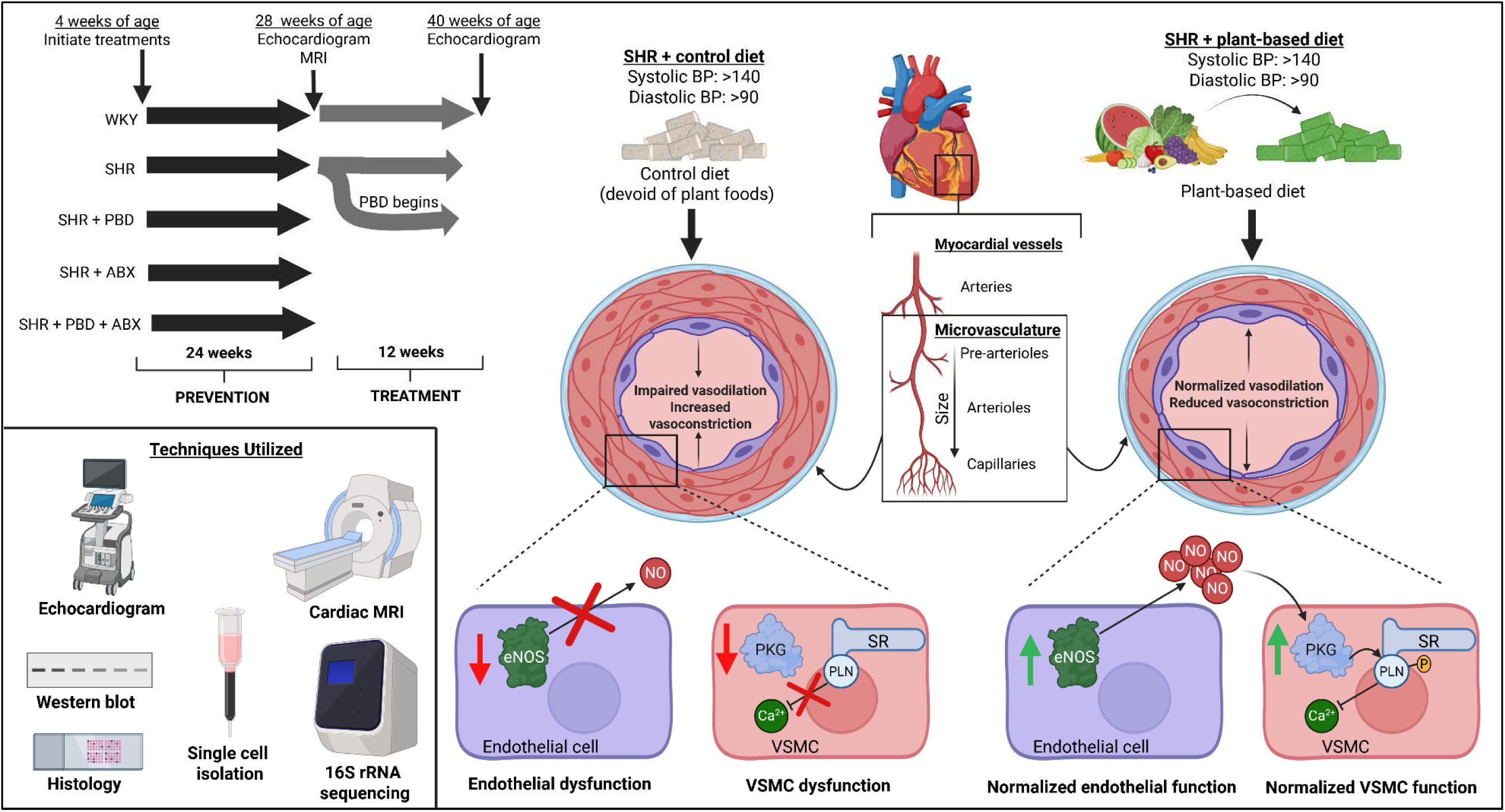

A plant-based diet prevented and reversed CMD without attenuating hypertension. Such amelioration of CMD was not negated by antibiotics and correlated with improved endothelial and VSMC function. Legend: ABX, antibiotics; BP, blood pressure; eNOS, endothelial nitric oxide synthase; NO, nitric oxide; PBD, plant-based diet; PKG, protein kinase G; PLN, phospholamban; SHR, spontaneously hypertensive rat; SR, sarcoplasmic reticulum; VSMC, vascular smooth muscle cell.

## 1. Introduction

Chest pain (angina pectoris) affects ∼112 million people globally and is classically associated with ischemic heart disease due to obstructive atherosclerosis^1^. Interestingly, up to two-thirds of women and one-third of men with angina who undergo coronary angiography are found to lack evidence of a blockage^2^. This condition is termed ischemia with non-obstructive coronary artery disease (INOCA)^3^, and is linked to increased risk of major adverse cardiovascular events, including myocardial infarction and heart failure^3^. Women are also more likely to have angina^4^, and are four times more likely than men to be re-hospitalized within six months following discharge^3^. An underlying pathological driver of INOCA is coronary microvascular dysfunction (CMD), an impairment of the cardiac microvessels, which are responsible for 80% of total vascular resistance^5^. CMD is diagnosed by a reduction in coronary flow reserve (CFR), the ratio of coronary blood flow at rest versus maximal blood flow. A major risk factor for CMD is hypertension, which is associated with reduced CFR^6^. CMD is associated with dysfunction of endothelial cells (ECs) and vascular smooth muscle cells (VSMCs)^7^. EC dysfunction largely results from reduced nitric oxide (NO) bioavailability due to downregulated expression of endothelial nitric oxide synthase (eNOS), which produces NO, and/or increased production of superoxide (O ^−−^), which reduces NO bioavailability via generation of peroxynitrite (ONOO^−^). VSMC dysfunction includes impaired vasodilation due to improper calcium handling via sarcoplasmic reticulum (SR) stress, and reactive oxygen species (ROS)-mediated inhibition of guanylyl cyclase, limiting protein kinase G activation (PKG)^8^.

CMD is treated by controlling cardiac risk factors, in combination with the use of anti-atherosclerotic, anti-anginal medications such as statins, angiotensin converting enzyme inhibitors, beta-blockers, calcium channel antagonists, and ranolazine for symptom management. However, CMD therapeutic strategies are not clearly defined, with patients having persistent symptoms and poor response to usual therapies in a subset^9^ ^5^. Furthermore, this multi-pharmacological approach is burdensome to patients and associated with a variety of adverse effects/intolerances and compliance issues. Thus, new approaches are urgently needed.

Consumption of diets rich in fruits and vegetables has long been known to be health-promoting, especially with respect to CVD prevention. Indeed, plant-based diets (PBDs) are associated with a reduced risk of ischemic heart disease^10^. This association appears unique to “healthy” PBDs^11^, characterized by increased intake of unprocessed plant-foods, such as whole fruits, vegetables, nuts, seeds, legumes, and whole grains, compared with “unhealthy” PBDs comprising processed foods (e.g., refined grains and added sugars)^11^. Some of the benefits PBDs stem from components not found on standard nutritional labels, including phytochemicals such as polyphenols, secondary metabolites of plants, which have beneficial molecular effects in attenuating CVD^12^. Once consumed, these phytochemicals undergo gut microbiota metabolism^13^, and these metabolites can exert their effects on target tissue. Indeed, in interventions that maximize the intake of unrefined plant-foods, reversal of atherosclerosis can be observed as well as improved myocardial perfusion in patients with coronary artery disease (CAD)^14–16^.

The extent to which PBD might mitigate CMD is largely unknown. Indeed, only a single published study to date used a dietary approach to attenuate CMD, namely calorie restriction, with poor efficacy^17^. Thus, we sought to examine the potential of a plant-based diet (PBD) in CMD. We developed approaches to measure CMD, which was indeed present in SHRs, and found a PBD prevented and reversed this disorder.

## 2. Methods

### Study Design

All animal use and procedures were approved by Georgia State University’s Institutional Animal Care and Use Committee (protocol #: A23025). Female animals were used to align with the fact that CMD afflicts women more severely than men^18–20^. Three-week-old, female, normotensive Wistar-Kyoto (WKY) rats and SHRs were purchased from Inotiv (West Lafayette, IN, USA) within the same respective in-bred colonies. Upon arrival, rats were doubly housed in an environmentally controlled animal care facility and maintained on 12-h light/dark cycles. Rats had free access to water and were maintained on a compositionally-defined open-source diet (Table S1 and Table S2; D23061303, Research Diets). After one week, SHRs either 1) continued the diet, 2) were placed on antibiotic (ABX) treatment to deplete the gut microbiota, 3) began the consumption of a PBD (Table S1 and Table S2; D23061304, Research Diets), or 4) consumed a PBD along with ABX treatment. WKY continued the control diet and did not consume ABX or PBD. To assess the potential of a PBD to treat CMD, a subset of SHRs on the control diet were switched to a PBD at 24 weeks of age and monitored for 12 additional weeks. A subset of WKY and SHR also continued the control diet for 12 weeks. Figure 1 outlines the animal study design.

**Figure 1.**
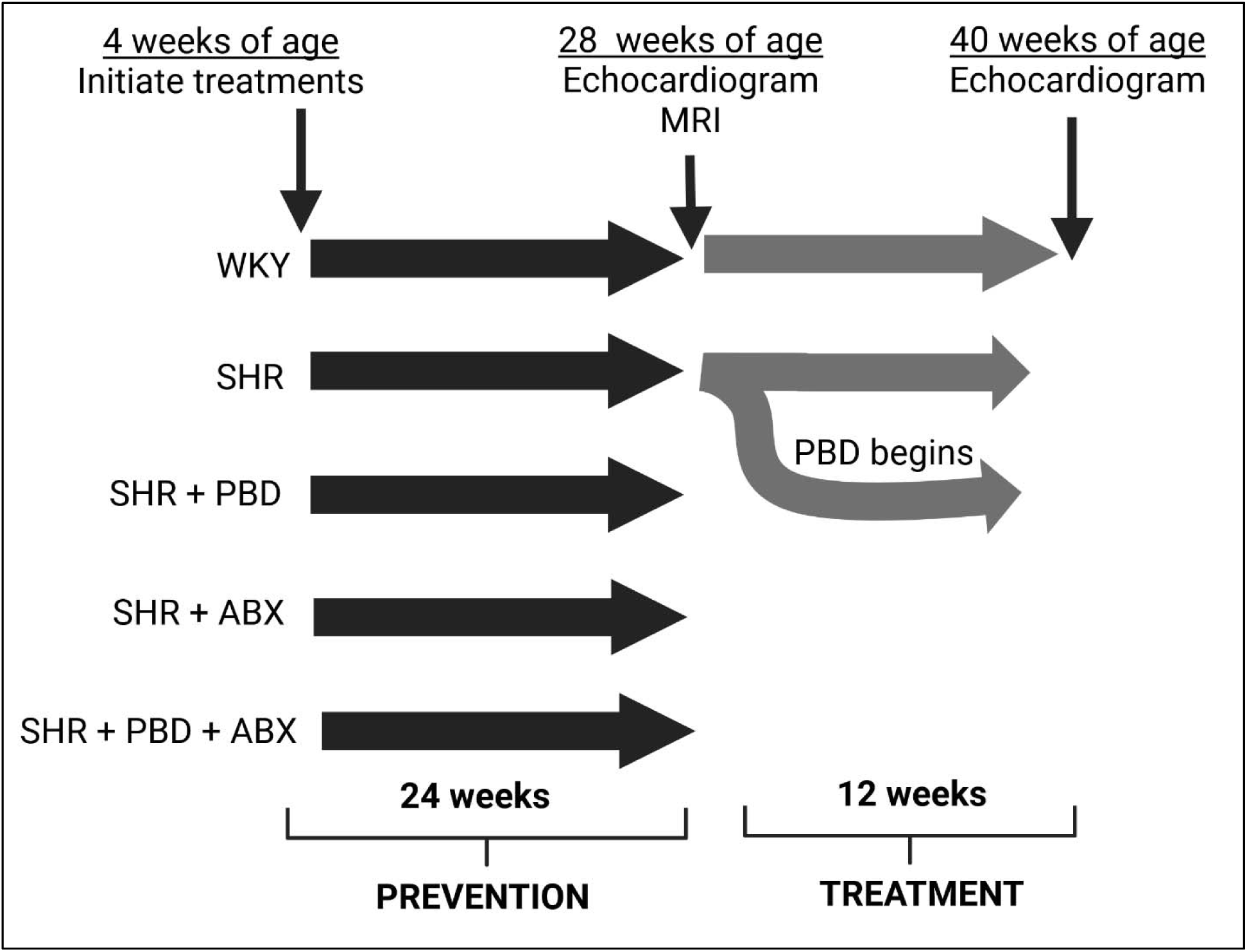
Animal study design.

The PBD comprised 28% (w/w) seven different plant foods: walnuts, black beans, red bell pepper, sweet potato, blueberries, Brussels sprouts, and lemon (4% each). These foods were selected for their high polyphenol content within their respective food categories^21^ (citrus, cruciferous vegetables, legumes, etc.) and their common consumption in the United States. Soy protein was used in place of casein in the PBD. The control diet was matched to the PBD in both macronutrient and micronutrient composition. Fiber was also matched, including in both soluble (inulin) and insoluble (cellulose) fiber (Table S1 and Table S2). Thus, the main known difference between the PBD and control diet was phytochemical content. The control diet did not contain any whole plant foods.

Regarding ABX treatments, the first 3 days of antibiotic treatment comprised of 1 g/L of ampicillin, neomycin sulphate, metronidazole, and 0.5 g/L of vancomycin in drinking water followed by a ⅓ concentration maintenance dose for the remainder of the study. Sucrose was added (0.5%) to increase palatability. This ABX water was changed weekly. Preliminary testing revealed that after 4 weeks of the maintenance ABX dose, 10^^−5^ fewer bacteria were in feces (∼4,602,842 bacteria/mg vs ∼57 bacteria/mg); thus, the ABX successfully reduced gut microbes.

### Body weight and food intake

The average body weight was obtained for each cage weekly, and food intake was also monitored weekly. To minimize oxidation of the PBD pellets, the PBD pellets were obtained from Research Diets in vacuum sealed 5 kg bags, and the portioned diet within the cages was discarded weekly. Because all diets were macronutrient-matched, dietary intake was reported as food intake in grams rather than energy intake.

### Fecal bacterial load and 16S rRNA sequencing

Animals were placed in autoclaved cages with autoclaved bedding, and feces were collected within 2 hours for microbial analysis. Feces were collected during weeks 2, 6, 12, 24, and 36. DNA was extracted and purified from frozen feces (80-120 mg) using DNeasy 96 PowerSoil Pro QIAcube HT kit, supplemented with PowerBead Pro Plates (Qiagen). The universal primers, 8F (5′-AGAGTTTGATCCTGGCTCAG-3′) and 338R (5′-CTGCTGCCTCCCGTAGGAGT-3′), were used to detect and quantify the amount of total fecal bacteria. Purified sample DNA was utilized for qPCR with Qiagen Quantinova SYBR green master mix (cat no. 208052) for reaction. A standard curve was utilized to predict the total number of bacteria per mg of feces. For 16s sequencing, V3-V4 region of 16S rRNA genes were amplified using the following primers: 341F (5′TCGTCGGCAGCGTCAGATGTGTATAAGAGACAGCCTACGGGNGGCWGCAG-3′) and 805R (5′GTCTCGTGGGCTCGGAGATGTGTATAAGAGACAGGACTACHVGGGTATCTAATCC-3′). PCR products of each sample were purified using Ampure XP magnetic purification beads. An index PCR was performed to attach dual barcodes and Illumina sequencing adapters using Nextera XT Index kit (Illumina). Final PCR products were verified on 1.5% DNA agarose gel, purified again using Ampure XP magnetic purification beads, and quantified using Pico dsDNA assay (Invitrogen). An equal molar of each sample was then combined as the library. The library was diluted and spiked with 5% PhiX control (Illumina) and sequenced by Illumina MiSeq Sequencing System (2 x 250bp). Demultiplexed fastq files were generated on instrument. Sequence reads were quality filtered by DADA2 plugin in Qiime2. Taxonomy was assigned using weighted Silva 138.1 classifier (animal distal gut). Raw 16s

### Blood pressure monitoring

Blood pressure was monitored every two weeks until 40 weeks of age via tail plethysmography using CODA high throughput non-invasive blood pressure system (Kent Scientific, Torrington, CT) in up to four rats simultaneously ^22^. All rats were encouraged to walk into the restraint which was adjusted to prevent excessive movement throughout BP recording. The occlusion cuffs were placed at the base of the tail and the volume pressure recording (VPR) cuffs were placed approximately 2 mm adjacent to the occlusion cuffs. Rats rested on the pre-heated heating platform for the duration of each experiment and tail temperatures remained between 35–37 °C. An additional heating blanket was used on top of the restraints for the first 10 minutes of acclimation to warm the animals sufficiently. BP experimental settings were as follows: occlusion cuffs were inflated to 250 mmHg followed by slow deflation over 15 s. The minimum volume changes, as sensed by the VPR cuff, was set to 20 μL. Each recording session consisted of 25 inflation and deflation cycles. During readings, the minimum accepted tail volume was 30 μL. Rats were habituated to the BP measurements over three timepoints before experimental recordings.

### Echocardiography: Cardiac function and coronary microvascular function

Echocardiography was performed during weeks 24, and 36 of the intervention. We have described our echocardiographic methods previously^23^. Vevo® 3100 Imaging Platform (Fujifilm Visual Sonics; Toronto, Canada) was used to measure functional and morphological parameters in M-mode with 4-wall left ventricle (LV) measurements. The probe frequency was 16 MHz. Animals were maintained on 2-3% isoflurane on a heated platform. The platform was longitudinally forward angled 5° so that the head was slightly elevated compared to the rest of the body. The ultrasound probe was positioned with a leftward lateral tilt of 60° to obtain the parasternal short axis view. The level of anesthesia was adjusted to obtain a target heart rate of 325 ± 25 beats per minute (bpm). In the prone angle, ophthalmic ointment was used on the eyes to prevent drying. A tourniquet was made using locking forceps and surgical porous tape at the base of the tail to dilate the lateral tail vein. A 26-gauge ¾ inch catheter (Cat #: 07-836-8494, Patterson Veterinary, Loveland, CO) was inserted into the lateral tail vein, the catheter was flushed with sterile PBS, and the catheter was capped to prevent bleeding. The animal was then flipped to the supine position, and hair clippers were used to shave the fur from the neckline to mid chest level. Residual body hair was removed with hair removal cream. Ultrasound transmission gel was applied to the contact area on the rat, and the LV was visualized, first in B-mode. M-mode was used once the largest part of the LV was visualized.

Following LV image capture, isoflurane was maintained at 2.5%, and the probe was moved cephalically until the aorta, right atrium, and right ventricular outflow tract were visualized, at which point color doppler was utilized to identify coronary flow. Scientific support from FujiFilm Visual Sonics assisted in the training of these techniques and obtainment of images ensuring proper identification of coronary flow, minimizing error in image precision. The mode was then changed to pulse wave, and the cursor was placed in the middle of the coronary flow, with minor adjustments to locate the point of maximal flow. The doppler angle was made parallel to the direction of flow, and this angle was not altered following baseline image capture. Coronary flow velocity is easily identifiable by the “sailboat” pattern of the velocity peaks. Following baseline coronary velocity capture, the catheter cap was removed, and a sterile 20-gauge 40-inch small bore extension (Cat #: 07-890-7617, Patterson Veterinary, Loveland, CO) was inserted into the catheter. This extension line was pre-filled with a sterile PBS solution containing adenosine at a concentration of 0.5 mg/mL which was attached to a filled 5 mL syringe on an infusion pump (Fusion 100-X Touch Pump, SAI Infusion Technologies, Lake Villa, IL). Infusion proceeded at a rate of 0.25 mg/kg BW per min for 6 min. Coronary flow velocity was monitored during this time, and maximal flow was captured. CFR was calculated as maximal velocity ÷ baseline velocity. An average of at least 2-3 consecutive beats were used for velocity measurements. While echocardiography has some subjectivity, the same operator was used for all measures ensuring consistency in results, and peak velocity was quantified during both baseline measures and maximal hyperemia measures. While in humans, 0.14 mg/kg BW per min of adenosine is typically used, our dosage was selected based on preliminary testing. It was found that maximal hyperemia was achieved with 0.25 mg/kg BW in rats without compromising cardiac function so severely that a hyperemic response was inhibited. Inhibited hyperemia was characterized by a heart rate that was too low and would not recover until infusion ceased.

### Cardiac magnetic resonance imaging (cMRI): cardiac microvascular endothelial function

Experiments involving cMRI were performed at the Advanced Translational Imaging Facility (ATIF) (Georgia State University, GA, USA). MRI Measurements were taken on week 24 of the intervention. Before commencing the scanning, animals placed in an induction chamber connected to an isoflurane anesthesia unit. The animals were induced with isoflurane (3-5% in oxygen). A tail vein catheter was placed on the rat for injection of L-NG-Nitroarginine methyl ester (L-NAME) after the baseline image collection. Then the rats were transferred to the MRI cradle and placed on an imaging platform (warmed to 37° C) where anesthesia was maintained with a nosecone (1-3% isoflurane) utilizing the SomnoSuite (Kent Scientific, Somnosuite, CT) small animal anesthesia delivery system. Upon induction of anesthesia, the animal’s eyes were lubricated with a sterile ophthalmic ointment. A thermocouple rectal probe was placed in the rectum to monitor core temperature. Respiratory rate was also monitored (SAI instruments Inc, NY), with a target rate of 40-60 breaths/min. Light paper tapes were used to minimize the movement of the animal body in addition to the stereotactic setup. To monitor cardiac rhythm, MRI compatible ECG gold disc electrodes were attached on the two front paws and right back paw using gel paste, and the other end of the leads were connected to an electro-optical converting box that converts the ECG signal for fiber optic transmission to the MRI scanner. Prior to commencing the MRI procedure, all four paws were assessed via toe pinch to assure an absence of a withdrawal response. MRI scanning was performed for ∼60 minutes using a Biospec Bruker 7T/21 cm MRI (Bruker Billerica, MA) inner diameter horizontal bore magnet using a specialized large 86 mm ^1^H transmit-receive volume radio frequency coil with an inner diameter of 72 mm.

The rat was scanned for structural mapping of heart before the tail-vein injection. A baseline localizer scan was obtained along the three orthogonal directions using the gradient echo scan (Gradient Recalled Echo). Then, a Bo map was obtained for the heart to correct any inhomogeneities in the magnetic field. Three additional reference scans are acquired, and they served as a reference to setup short axis or four chamber views. Using the localizer, first an axial scan with 6 slices of 0.8 mm thickness of the rat heart was acquired using GRE scan. From the previous axial slices, coronal slices of rat heart (perpendicular to axial) were imaged with similar parameters as above. Additional scans of sagittal slices from the coronal scan through the outflow tract of the left ventricle and the apex were taken. From these sagittal images, a short axis view (axial images) of rat heart using CINE IG-FLASH (intragate FLASH MRI sequence, retrospectively triggered). This allows the cardiac imaging of the rat heart without use of ECG signals and completely free of motion artifacts. One min scans with flip angles of 25° with TR =21.3 msec, TE =1.9 msec, matrix size of 128 x 128, FOV of 50 x 50 mm, oversampling of 30, three movie frames were used for scans.

Coronary microvascular endothelial dysfunction was assessed via MRI by utilizing L-NAME, an inhibitor of nitric oxide synthesis ^24^. L-NAME was prepared in phosphate buffered saline and provided to animals at a concentration of 75 mg/kg of BW ^25^. The L-NAME solution was sterilized with 0.22 m PVDF membrane filter prior to tail-vein injection via a catheter. After the injection, the same scanning was repeated at minutes: 4, 8, 12, 16, 20 and 24. This was assessed semi-quantitatively using MATLAB (R2024a, Mathworks Inc) with the medical imaging quantitative software, Aedes 1.0 (University of Eastern Finland, Finland). For analysis, a region of interest (ROI) was drawn along the myocardium to segment the endomyocardial borders in the end-systolic and end-diastolic frames using the multi-slice short axis scan images, and the signal intensity values from each ROI were used for volumetric measurements at various time points. All ROIs were normalized to each animal’s respective baseline value, as raw ROI values cannot be compared between animals due to basal differences in fluid dynamics.

### Cell isolation and culture

Rats were euthanized with CO_2_ followed by decapitation with guillotine. Under aseptic conditions in a laminar flow chamber, the chest of the rat was sprayed with 70% ethanol. The chest was then opened with autoclaved surgical instruments, the heart was excised, atriums and aorta removed, and the remaining ventricles were placed in ice-cold HBSS (21-023-CV, Corning). The heart was transferred to 10 mL of VSMC growth medium (R311K-500, Cell Applications) in a 60 mm dish with 1 mg/mL of sterile-filtered DNAse 1 (LS002140, Worthington) and collagenase 2 (LS004176, Worthington), as well as 1 U/mL dispase 2 (17105041, ThermoFisher Scientific) for digestion. The heart was cut into small pieces with surgical scissors, transferred to a 15 mL tube with the digestion medium, then gently rocked for 30 minutes at 37° C. After which, a metal, luer lock 16-gauge blunt needle (14-815-604, Fisher Scientific) was used with a 35 mL syringe to mechanically disrupt the tissue. Care was taken to not exert intense pressure as to not kill the cells. Tissue digestion continued for another 30 min., followed by filtering through a 70 µm cell strainer. Cells were centrifuged at 300 g, and tissue debris was removed with debris removal solution (130-109-398, Miltenyi Biotec) per the manufacturer protocol. Red blood cells were then lysed and removed per the manufacturers protocol (130-094-183, Miltenyi Biotec).

Following centrifugation, cells were resuspended in CD45-conjugated magnetic microbeads (130-109-682, Miltenyi Biotec) and filtered through a LD column (130-042-901, Miltenyi Biotec) in a magnetic field. The filtrate was collected, centrifuged, then seeded in a 6-well plate (two wells) with complete VSMC growth medium, while the CD45-labeled cells were collected and lysed in radioimmunoprecipitation assay (RIPA) buffer with protease and phosphatase inhibitors. Cultured cells were allowed to become confluent (5-7 days) with medium change every other day. Following confluence, cells were placed in starvation medium (10% of the growth supplement; 0.5% FBS) for 24 h. Cells were then lysed at passage 0 in supplemented RIPA or treated with or without freshly prepared sodium nitroprusside (100 nM) for 5 min to induce nitric oxide (NO)-mediated signaling. Alternatively, cells were collected to determine population type with flow cytometry.

### Flow cytometry

For flow cytometry analysis, the prepared cells were blocked with 2.4G2 (anti-CD16/anti-CD32) at concentration 1 μg/million cells in 200μl PBS for 15 minutes at 4°C, followed by a wash with PBS twice to remove all residue of 2.4G2. Cells were then incubated with isotypes IgG1 kappa eF660 (Cat #: 50-4301-82) and IgG2a kappa isotype AF488 (Cat #: 53-4321-80) or a cocktail of conjugated monoclonal antibodies CD31 (25-0310-82, ThermoFisher Scientific), CD90 (12-0900-81, ThermoFisher Scientific), and α-SMA (ab202295, Abcam) at 4°C for 30 minutes in dark. Multi-parameter analysis was performed on a CytoFlex (Beckman Coulter) and analyzed with FlowJo software (Tree Star).

### Protein analysis

In addition to cell lysates, the hypothalamus and heart were also excised from animals for protein analysis. Immediately after sacrifice, the skull was cut open with bone scissors, and the brain was detached from the cranium. Using small, blunt forceps to hold the brain in place, a curved spatula was placed under the hypothalamus and gently pushed upwards to detach the hypothalamus. The hypothalamus and whole hearts were immediately frozen in liquid nitrogen and stored at −80° C. Hypothalamus was homogenized in 400 µL supplemented RIPA using a glass Dounce homogenizer. LV tissue from the heart (20-40) mg of was also homogenized in the same way. All protein lysates from cells and tissue were centrifuged at 16,000 x g for 20 min and supernatants were collected. Alternatively, nuclear extraction was performed on the LV utilizing a commercially available kit (78833, ThermoFisher Scientific) as well as glutathione isolation (703002, Cayman Chemical). Protein concentration of lysates was determined using the DC protein assay kit (BioRad Laboratories). For glutathione quantification of the LV, 5 µg of protein was used in each well of the assay. For Western blot, 60 µg of protein from tissue or 20 µg from cell lysates were separated in 8-15% SDS-PAGE gels and transferred to polyvinylidene difluoride (PVDF) membranes using Trans-Blot Turbo (BioRad Laboratories). Enhanced chemiluminescence (WBLUF0500, Millipore Sigma) was used to determine expression of proteins involved in redox, inflammation, and signaling depending on the tissue/cell of interest. The density of protein bands was quantified using Image Lab 6.0 (BioRad Laboratories, Hercules, CA) which was normalized to total lane protein. Phosphorylated proteins were normalized to their respective total protein. If phosphorylated and total proteins were on different membranes, then each protein was first normalized to the total lane protein followed by normalization of phosphorylated protein to total protein from each respective membrane.

From Cell Signaling Technologies, antibodies include phospho-NF-κB (3033), NF-κB (4764), c-Jun (60A8), phospho-c-Jun (9261), phospho-p38MAPK (4511), p38MAPK (8690), phospho-SAPK/JNK (4668), SAPK/JNK (9252), SOD2 (13141), SOD1 (37385), nitrotyrosine (92212), phospholamban (14562), p22phox (27297), p47phox (63290), eNOS (32027), phospho-eNOS (9571), VASP (3112), phospho-VASP (3114), Rabbit secondary (7074), and Mouse secondary (7076). From Abcam, antibodies include phospho-phospholamban (ab308132). From R&D systems, antibodies include GPx1 (AF3798), 4-HNE (MAB3249), and goat secondary (HAF109).

### Histological analysis

Following sacrifice, hearts were stored in 10% neutral-buffered formalin for 24 h, then moved to 70% ethanol. Tissue samples were dehydrated, embedded in paraffin, and used for histological analysis. Coronal cuts at 10 µm thickness were made. Hematoxylin and eosin (H&E) staining was used to assess morphology, and Sirius red staining was used to assess the extent of fibrosis. Fibrosis was assessed with ImageJ, using the Fiji software plugin. 10x images of areas with peak fibrosis were used for quantification. Blood vessels and collagen attachment sites were avoided for the analysis.

### Serum nitric oxide metabolites

Following decapitation, blood was collected, allowed to clot for 40 min. at room temperature, then spun at 500 g for 10 min. The serum was collected and used for the measurement of nitric oxide metabolites using a commercially available fluorometric kit (780051, Cayman Chemical).

### Statistical analysis

GraphPad Prism (San Diego, CA) was used for all statistical analyses. Statistical tests depended on the assessment and hypotheses. Statistical tests are stated in the figure legends. These include one-way ANOVA followed by Dunnett’s post-hoc analysis, two-way ANOVA, or Student’s t-test. Values are represented as mean ± standard deviation (SD). Data were deemed significant if P ≤ 0.05. Sample size determinations were made based on prior experience in conducting measures of interest.

## 3. Results

### PBD did not ameliorate hypertension or impact cardiac function in SHRs

Tracking of body weight indicated that, in accord with other studies^26^, SHRs exhibited moderately reduced weight gain relative to age-matched WKYs (Figure 2A & B). This difference was accompanied by SHRs exhibiting a slight reduction in cumulative food intake during this period (Figure 1C & D). PBD and/or ABX did not significantly impact weight at 28 weeks, when heart function was to be assessed. Final body weight at week 28 and 40 weeks of age is provided in Table S3.

**Figure 2.**
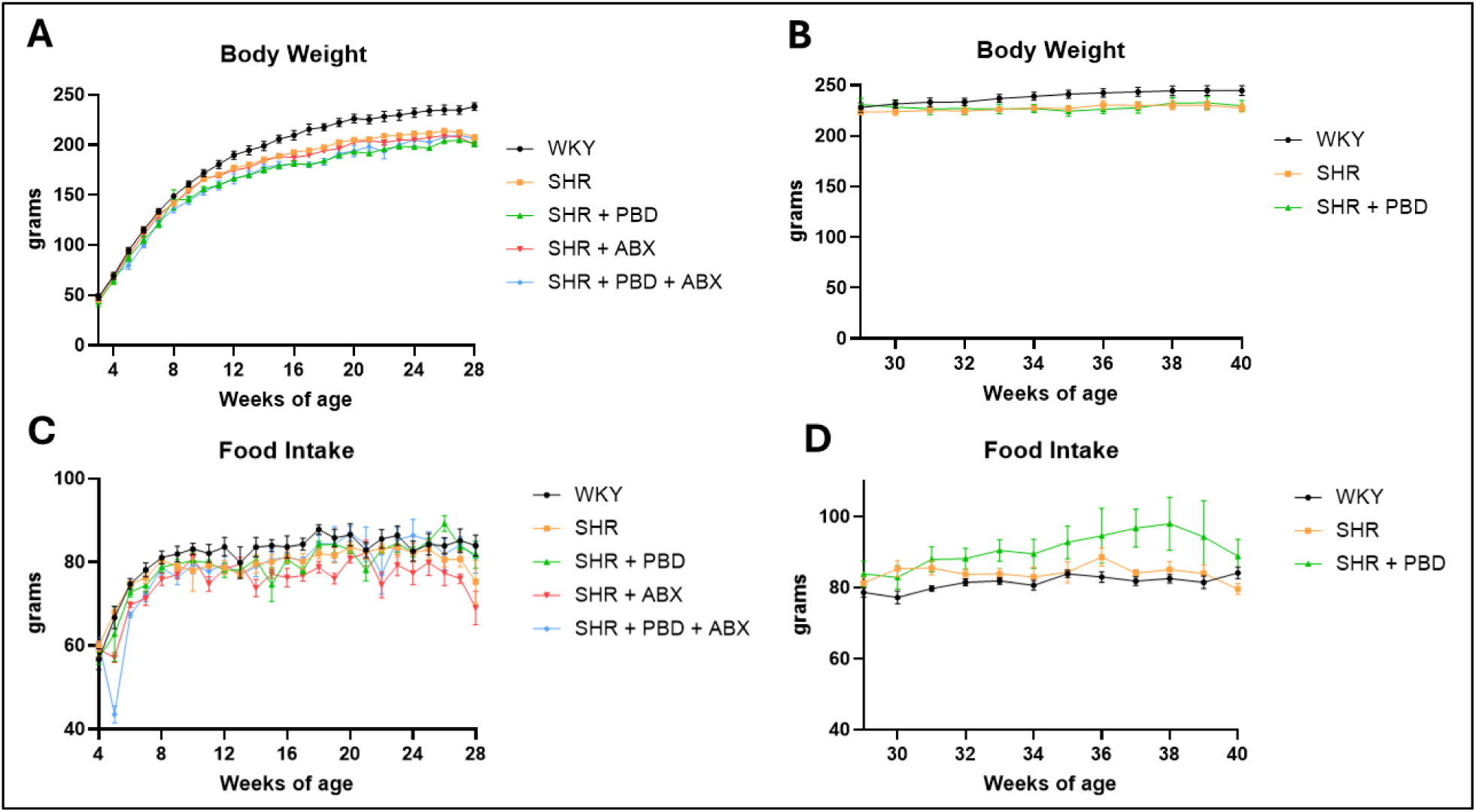
Body weight and food intake. Wistar-Kyoto (WKY) rats or spontaneously hypertensive rats (SHR) consumed a purified diet or 28% (w/w) plant-based diet (PBD) for 24 weeks with or without antibiotics (ABX). After 24 weeks, a subset of SHRs were switched from the control diet to PBD, while a subset of SHRs and WKY continued the control diet for 12 additional weeks. (A, B) Body weight and (C, D) food intake were measured weekly, and the average of both animals per cage was used (n = 8– 14/group). Data are expressed as mean ± SD.

The assessment of microbiota composition via 16S rRNA sequencing indicated that, as expected, microbiomes of SHRs randomized to receive the control or PBD were similar at baseline (Figure 3A). However, microbiomes of rats who were fed the control and PBD clearly differed from each other at all timepoints assessed following diet initiation (Figure 3A & B; Figure S1). Despite this, these changes did not significantly impact alpha diversity between groups (Figure S2). A similar pattern of PBD-induced changes in the microbiome was observed for animals that began consuming PBD at 24 weeks of age (Figure 3B). PBD-induced changes in the microbiome included changes in several genera whose abundance was significantly (adjusted p-value<0.01) enriched (blue) or depleted (orange) following PBD invitation at weeks 24 or 36 (Figure 3C & D).

**Figure 3.**
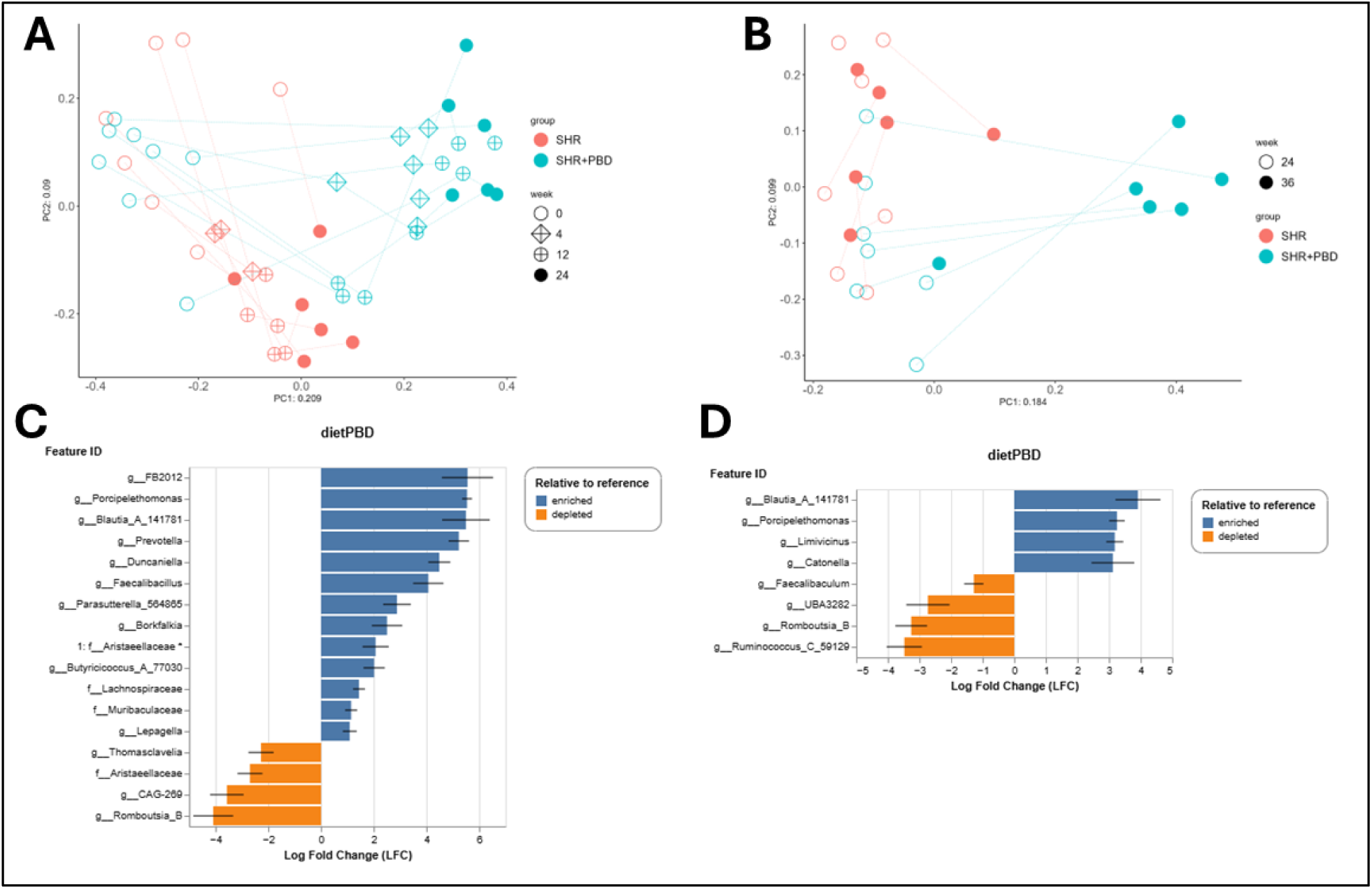
Gut microbiota beta diversity and bacterial taxa of differential abundance in SHRs consuming a control diet or PBD. Spontaneously hypertensive rats (SHR) consumed a purified diet or 28% (w/w) plant-based diet (PBD). After 24 weeks, a subset of SHRs were switched from the control diet to PBD, while a subset of SHRs continued the control diet for 12 additional weeks. Feces were collected at weeks 0, 4, 12, 24, and 36 for 16S rRNA sequencing. (A, B) PCoA plot of Bray-Curtis distance and (C, D) enrichment or depletion of bacterial genus after PBD diet is illustrated. Panels A and C reflect weeks 0-24 during the prevention phase, while panels B and D reflect weeks 24-36 during the treatment phase.

Blood pressure measurements revealed that, as expected, WKY exhibited normal blood pressure while SHRs had significantly increased systolic and diastolic blood pressure. Neither PBD nor antibiotics ameliorated hypertension (Figure 4). Final blood pressure values at weeks 28 and 40 weeks of age are provided in Table S4.

**Figure 4.**
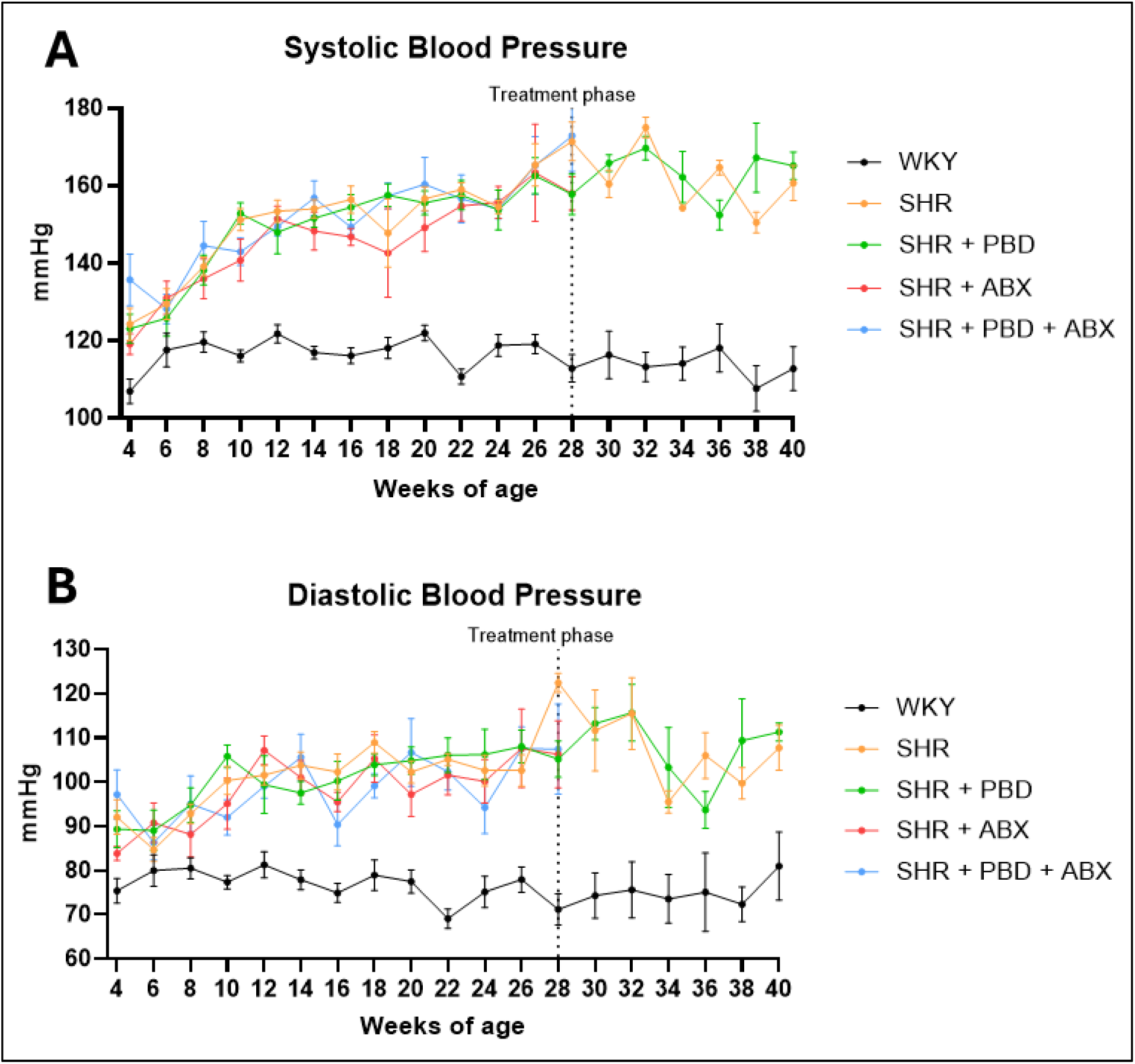
Blood pressure over time. Wistar-Kyoto (WKY) rats or spontaneously hypertensive rats (SHR) consumed a purified diet or 28% (w/w) plant-based diet (PBD) for 24 weeks with or without antibiotics (ABX). After 24 weeks, a subset of SHRs were switched from the control diet to PBD, while a subset of SHRs and WKY continued the control diet for 12 additional weeks. (A) Systolic blood pressure and (B) diastolic blood pressure was measured every two weeks with tail plethysmography (*n* = 5–14/group). Data are expressed as mean ± SEM.

### A plant-based diet prevented CMD development and reversed established CMD

Cardiac function of our cohort of 28-week-old female SHRs was assessed by measuring ejection fraction (EF) via echocardiography (Figure 5A, B). While EF values of female SHRs did not fall within the dysfunctional range, a result expected for their age^27^, they were nevertheless significantly reduced compared to age-matched WKYs. Consistent with the absence of marked cardiac dysfunction, there was also no significant difference in LV mass normalized to body weight among WKYs and SHRs (Figure 5A, C). Nonetheless, at week 24, both LV internal diameter during systole (LVIDs) and during diastole (LVIDd) was significantly increased in SHRs compared to WKY (Figure 5A, D, E), indicating LV dilation. None of these measures in SHRs were impacted by PBD, ABX, or the combination thereof, nor did PBD impact these parameters during the treatment phase of the study (Figure 6).

**Figure 5.**
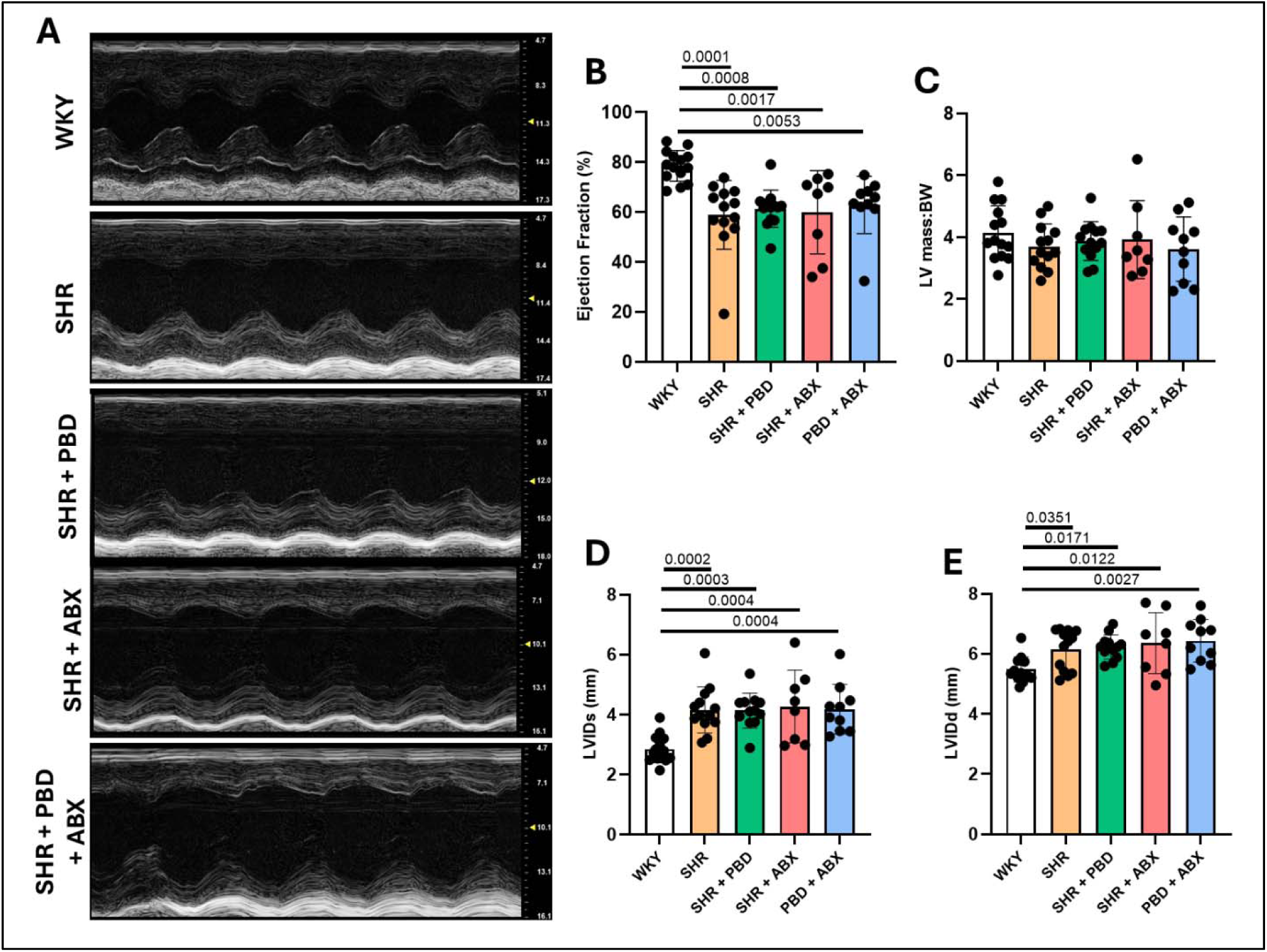
Cardiac function and morphological characteristics during the prevention phase (week 24). Wistar-Kyoto (WKY) rats or spontaneously hypertensive rats (SHR) consumed a purified diet or 28% (w/w) plant-based diet (PBD) for 24 weeks with or without antibiotics (ABX). (A) Cardiac function was assessed by (B) ejection fraction (EF) with echocardiogram. (C) Left ventricle (LV) morphology was assessed with LV mass normalized to body weight (BW), as well as (D) LV internal diameter during systole (LVIDs) and (E) diastole (LVIDd). Data were statistically assessed with one-way ANOVA and Dunnett’s post-hoc test (n = 8–14/group). Significance (P ≤ 0.05) is denoted in the graphs with exact *P*-value. Data are expressed as mean ± SD.

**Figure 6.**
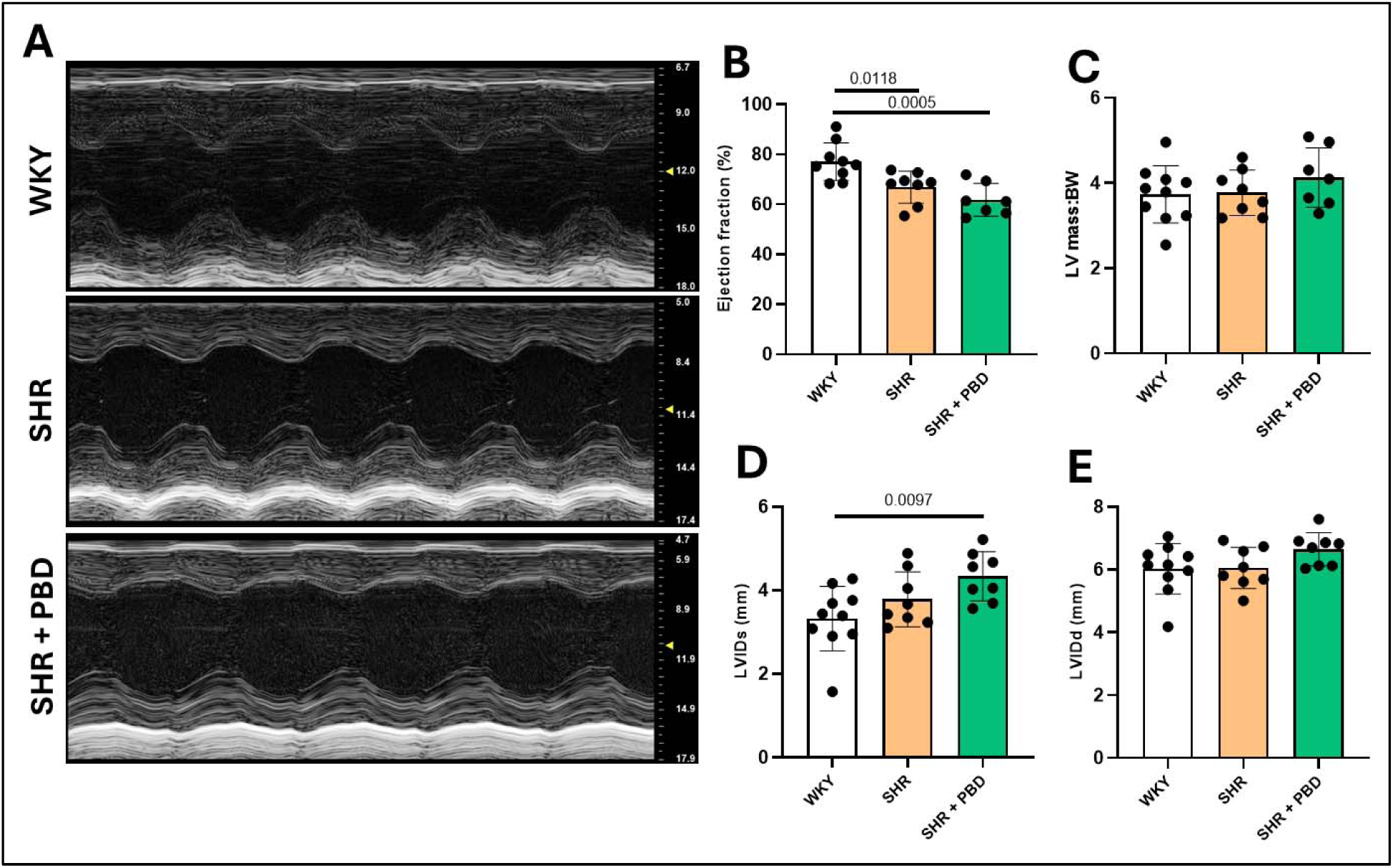
Cardiac function and morphological characteristics during the treatment phase (week 36). Wistar-Kyoto (WKY) rats and spontaneously hypertensive rats (SHR) consumed a purified diet for 24 weeks. After 24 weeks, a subset of SHRs were switched from the control diet to PBD, while a subset of SHRs and WKY continued the control diet for 12 additional weeks. (A) Cardiac function was assessed by (B) ejection fraction (EF) with echocardiogram. (C) Left ventricle (LV) morphology was assessed with LV mass normalized to body weight (BW), as well as (D) LV internal diameter during systole (LVIDs) and (E) diastole (LVIDd). Data were statistically assessed with one-way ANOVA and Dunnett’s post-hoc test (n = 7–9/group). Significance (P ≤ 0.05) is denoted in the graphs with exact *P*-value. Data are expressed as mean ± SD.

The hallmark of CMD is a reduction in CFR, which we measured by echocardiography. While a normal range of CFR in rats has not been well established, CFR levels were indeed lower in SHRs than WKYs (CFR of WKYs and SHRs was 2.25 ± 0.38 vs. 1.70 ± 0.39, respectively p-0.01) supporting our presumption that SHRs indeed develop CMD (Figure 7A, B). Most importantly, PBD significantly increased CFR, restoring it to the levels of WKYs (CFR of PBD-fed SHRs was 2.32 ± 0.54, p=0.01 for SHRs fed control vs PBD diet) supporting our hypothesis that PBD could prevent CMD. The extent to which gut microbiota contributed to the benefits of PBD was less clear. Specifically, CFR of control SHRs with ABX did not differ significantly from WKY or control SHRs without ABX. Thus, determining the role of the gut microbiota in this model will require future studies.

**Figure 7.**
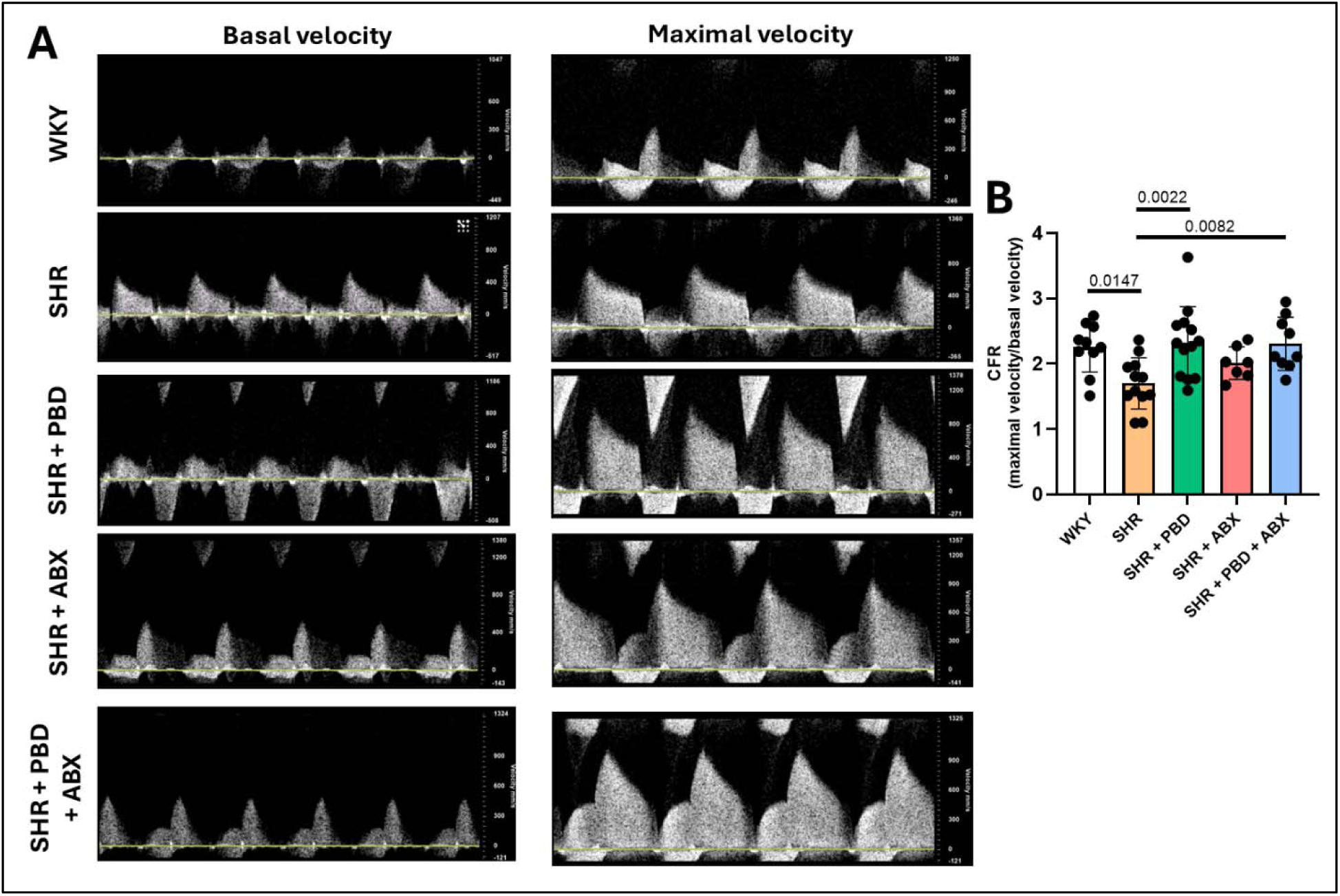
Coronary microvascular function during the prevention phase (week 24). Wistar-Kyoto (WKY) rats or spontaneously hypertensive rats (SHR) consumed a purified diet or 28% (w/w) plant-based diet (PBD) for 24 weeks with or without antibiotics (ABX). Coronary flow reserve (CFR) was calculated as maximal velocity prior to adenosine infusion ÷ basal velocity during adenosine infusion (0.25 mg/kg/min). Data were analyzed with one-way ANOVA and Dunnett’s post-hoc test (n = 7–13/group). Significance (P ≤ 0.05) is denoted in the graphs with exact *P*-value. Data are expressed as mean ± SD.

Echocardiograms performed on the second cohort of SHRs in the treatment study showed low and similar levels of CFR, at 24 weeks, in groups of animals that were to remain on the control diet or be switched to PBD. In accordance with the notion that hypertension-induced CMD continues to worsen with time, mean CFR levels declined between 24 and 36 weeks in SHRs fed the control diet in 5 of 8 SHRs assessed (Figure 8A). In contrast, CFR levels increased over this period in PBD-fed SHRs resulting in significantly higher CFR level in PBD-fed SHRs compared to SHRs fed the control diet (Figure 8B, C). The increase in CFR following PBD consumption resulted in CFR levels that were similar to WKYs, indicating that PBD had reversed CMD (Figure 8D, E).

**Figure 8.**
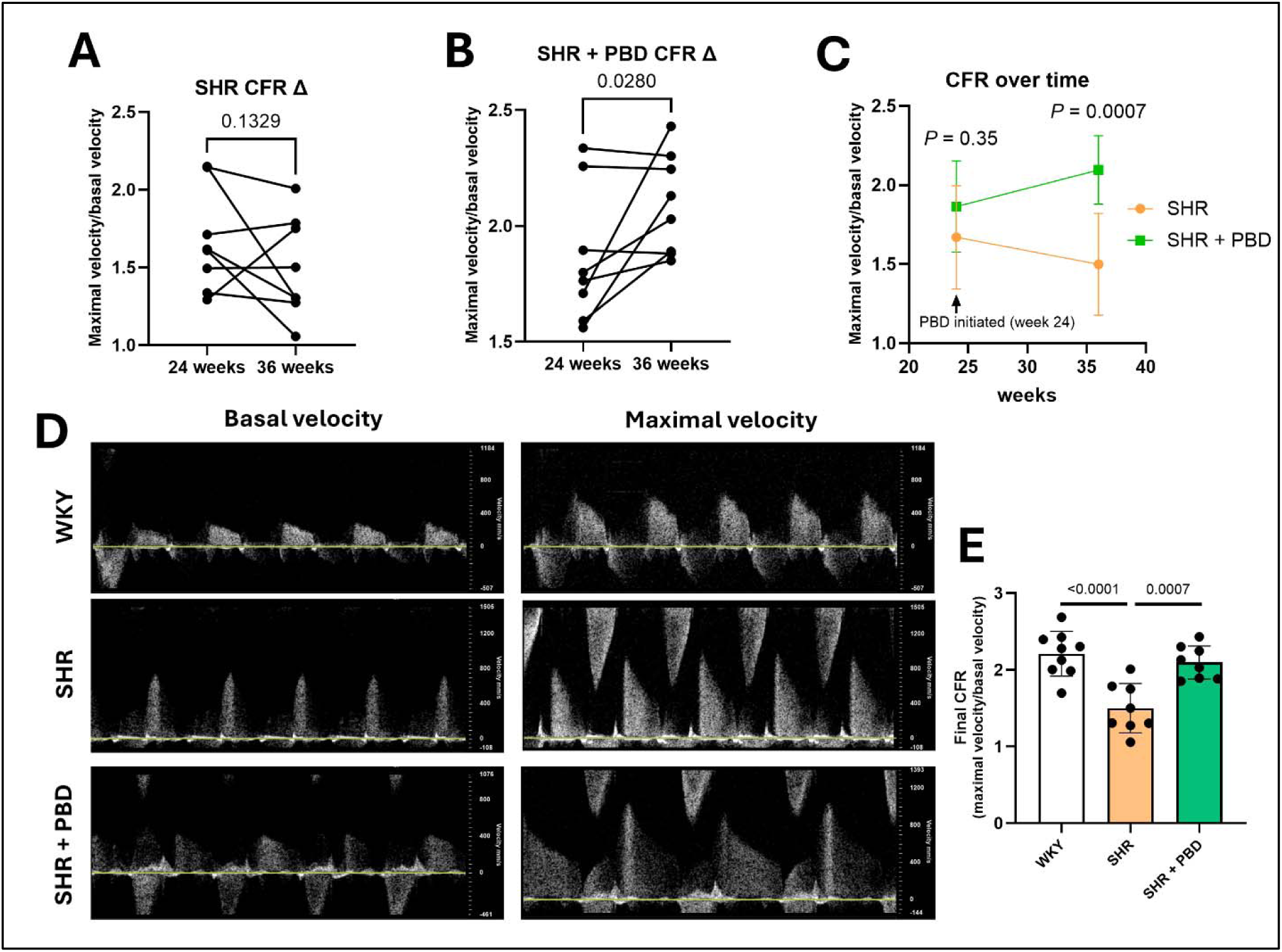
Coronary microvascular function during the treatment phase (week 36). Wistar-Kyoto (WKY) rats and spontaneously hypertensive rats (SHR) consumed a purified diet for 24 weeks. After 24 weeks, a subset of SHRs were switched from the control diet to PBD, while a subset of SHRs and WKY continued the control diet for 12 additional weeks. Coronary flow reserve (CFR) was calculated as maximal velocity prior to adenosine infusion ÷ basal velocity during adenosine infusion (0.25 mg/kg/min). Data was statistically assessed with (A, B) Student’s paired t-test, (C) two-way ANOVA with Šídák’s multiple comparisons test, or (E), one-way ANOVA with Dunnett’s post-hoc test (n = 8–9/group). Significance (P ≤ 0.05) is denoted in the graphs with exact *P*-value. Data are expressed as mean ± SD.

### A plant-based diet treats CMD in an endothelial-dependent and -independent manner

To investigate the cellular mechanisms by which PBD consumption prevented CMD, cardiac magnetic resonance imaging (cMRI) was employed (Figure 9). We evaluated myocardial blood flow (MBF) before and after the intravenous injection of 75 mg/kg BW L-NG-Nitroarginine methyl ester (L-NAME), an eNOS inhibitor. The rationale for this approach was that if eNOS was functioning normally, then L-NAME would dramatically reduce MBF. If eNOS was impaired, the provision of L-NAME would have little impact in reducing MBF. This was assessed by observing intensity changes of the myocardium (Figure 9A). Indeed, L-NAME injection resulted in a visually appreciable reduction in myocardial blood flow by 8 min in WKYs but not SHRs (Figure 9B). Image analysis-based quantitation confirmed this, as WKYs exhibited a stark drop in MBF in response to L-NAME injection while injection of PBS did not impact this parameter, indicating the experimental approach was working as expected (Figure 9C, D). In contrast, SHRs consuming the control diet exhibited only a modest reduction in MBF in response to L-NAME, indicating endothelial dysfunction. PBD-fed SHRS exhibited L-NAME-induced reductions in MBF that were at least as robust as that of WKYs indicating that PBD had fully prevented hypertension-induced cardiac microvascular endothelial dysfunction. ABX did not alter L-NAME induced changes in MBF in control- or PBD-fed SHRs, arguing that this aspect of PBD’s benefit was not microbiota-mediated. The preservation of eNOS activity in SHRs consuming PBD was associated with increased levels of NO metabolites, irrespective of ABX (Figure S3).

**Figure 9.**
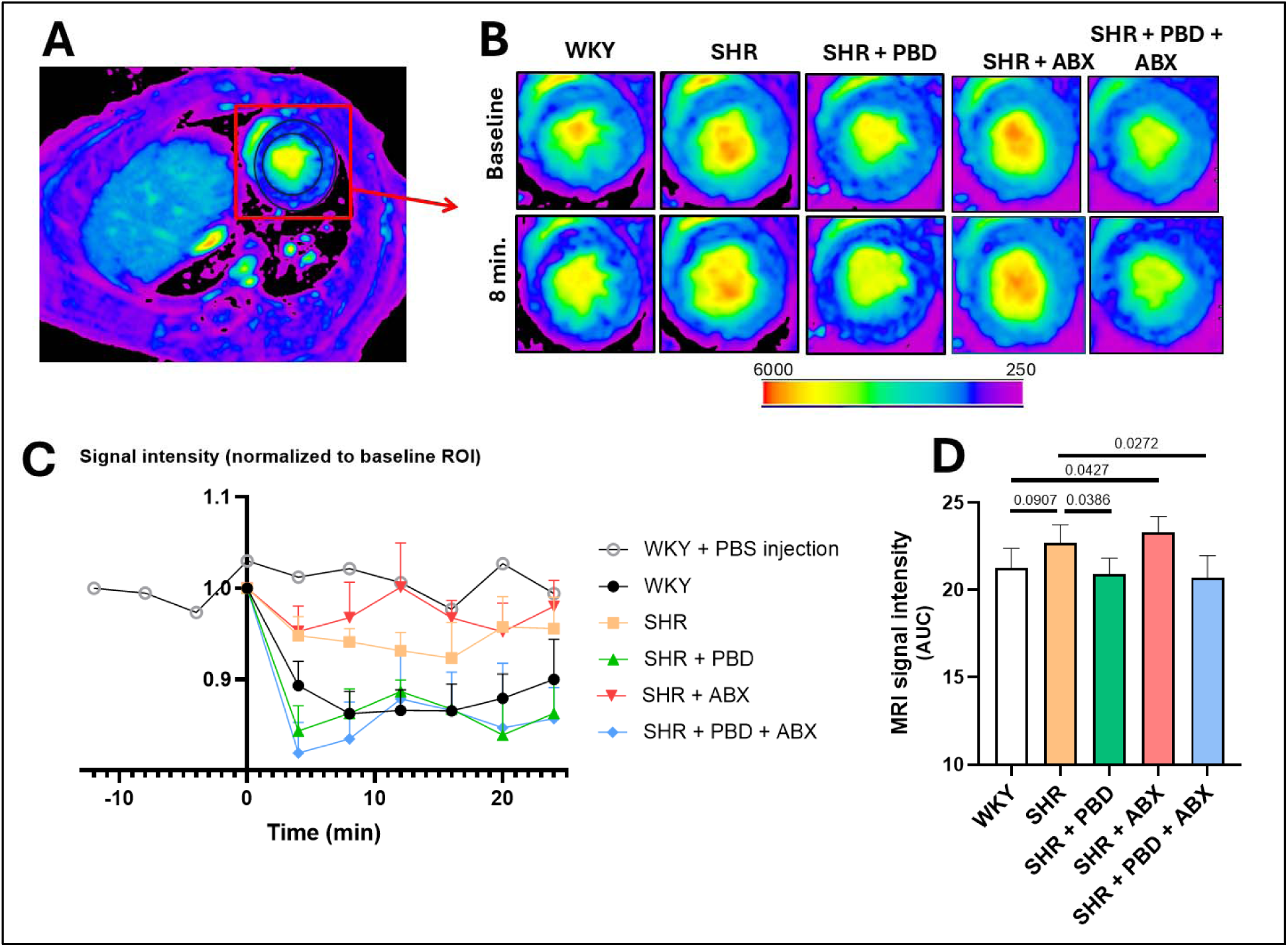
Coronary microvascular endothelial function via cMRI. Wistar-Kyoto (WKY) rats or spontaneously hypertensive rats (SHR) consumed a purified diet or 28% (w/w) plant-based diet (PBD) for 24 weeks with or without antibiotics (ABX). (A) The region of interest (ROI) of the myocardium used for analysis is outlined in black. A baseline MRI scan was acquired, followed by L-NAME injection, and subsequent scans every 4 min for a total of 24 min. In addition, one WKY was scanned every 4 min for 12 min prior to PBS injection, and every 4 min for 24 min after injection. (B) Representative cardiac MRI images are provided for baseline and 8 min. (C) Signal intensity over time, (D) which was statistically quantified with AUC analysis followed by one-way ANOVA and Dunnett’s post-hoc analysis (n = 4–6/group). Significance (P ≤ 0.05) is denoted in the graphs with exact *P*-value. Data are expressed as (C) mean ± SEM or (D) mean ± SD.

eNOS is a critical mediator of endothelial function and thus might have played a role in mediating the beneficial impacts of PBD. To probe this notion, we assessed LV eNOS and phospho-eNOS (Ser^1177^) levels via western blot in animals from the treatment study. We found that, PBD-fed SHRs exhibited elevated eNOS and phospho-eNOS relative to SHRs that consumed the control diet (Figure 10A–C). We envisaged that PBD’s potentiation of eNOS might impact VSMCs. Hence, we measured ventricular VSMC expression of phospho-vasodilator-stimulated phosphoprotein (VASP), which is downstream of PKG, activated by NO signaling, in cells treated or not, with sodium nitroprusside (SNP), a compound that releases NO. Phospho-VASP expression was markedly increased following SNP treatment compared to untreated cells (Figure S4), supporting the role of SNP in NO-induced signaling. Relative to WKYs, SHRs exhibited a marked decrease in NO-induced phospho-VASP levels, while PBD treatment fully restored phospho-VASP (Figure 10D, E). Analysis of the cell prep on which this assay was performed revealed both VSMCs (54.3%) and fibroblasts (45%) (Figure S5). VASP phosphorylation also occurs in fibroblasts. Thus, to validate NO signaling in VSMCs, phosphorylation of the VSMC-specific sarcoplasmic reticulum marker, phospholamban (PLN), a downstream target of PKG resulting from NO activity, was assessed (absent in fibroblasts)^28^. Phosphorylation of PLN results in the sequestration of Ca^2+^, facilitating VSMC relaxation. Mirroring phospho-VASP, phospho-PLN in VSMC was significantly increased in SHRs consuming PBD compared to SHRs consuming the control diet (Figure 10D, F). These benefits may have been localized to microvascular circulation. For example. isolation of aortic and mesenteric artery protein revealed that eNOS expression was diminished in SHRs regardless of diet; suggesting that conduit blood vessels were impaired, potentially driving the hypertension (Figure S6). However, aside from the microvasculature of the heart, we also found that eNOS protein expression in microvasculature of kidney and liver tissue was also diminished in SHRs, a detrimental effect ameliorated by PBD supplementation (Figure S7). Thus, a PBD exerted distinct effects between conduit vessels and microvasculature.

**Figure 10.**
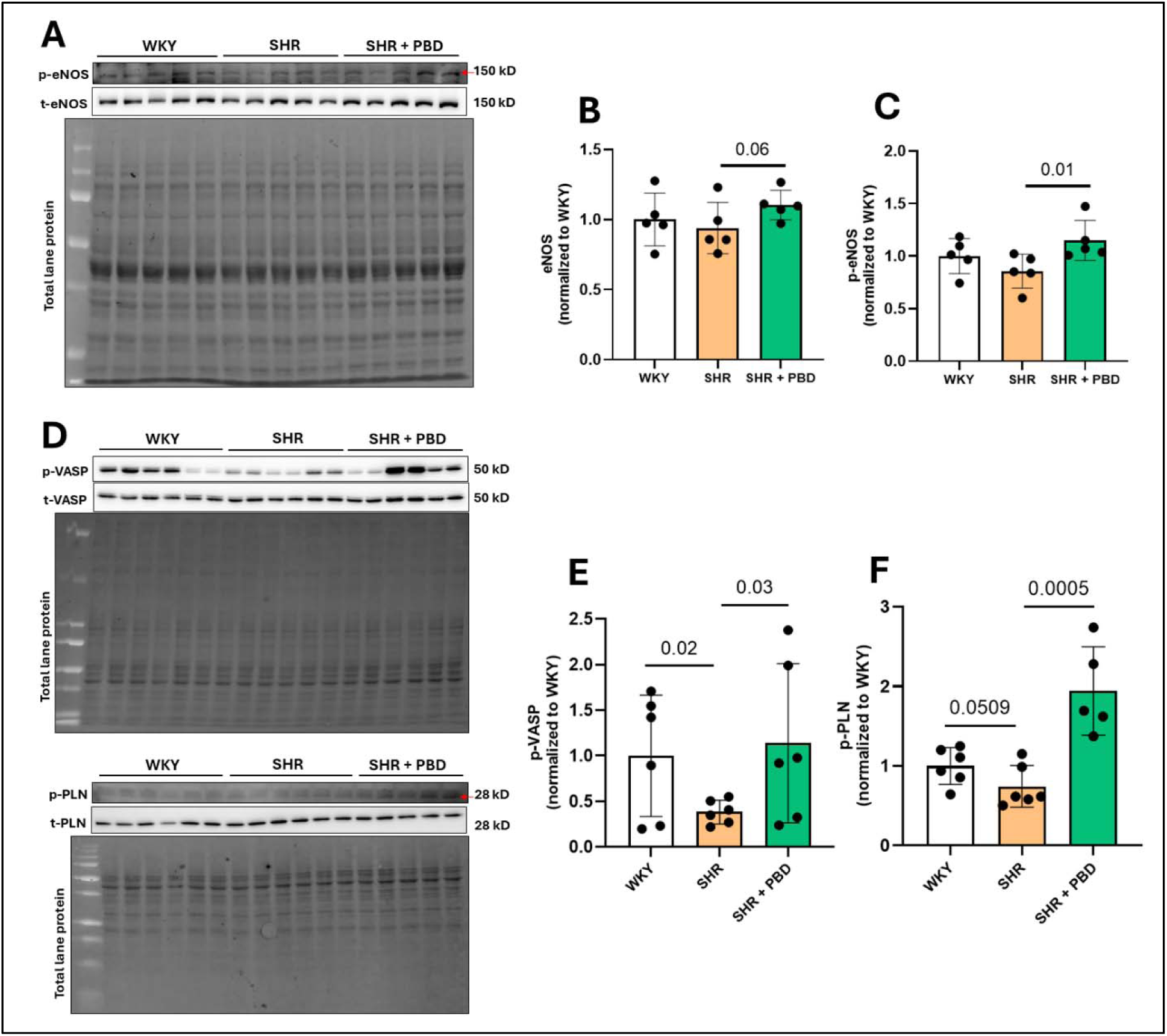
Cardiac endothelial- and VSMC-dependent cellular mechanisms. Wistar-Kyoto (WKY) rats and spontaneously hypertensive rats (SHR) consumed a purified diet for 24 weeks. After 24 weeks, a subset of SHRs were switched from the control diet to PBD, while a subset of SHRs and WKY continued the control diet for 12 additional weeks. (A) Left ventricular protein was isolated and (B) endothelial nitric oxide synthase (eNOS), as well as phospho-eNOS (C) was probed via western blot. Vascular smooth muscle cell (VSMC)-comprised cells were isolated from the heart. Following culture and confluence, cells were placed in starvation medium (10% of the growth supplement; 0.5% FBS) for 24 h. Cells were then lysed at passage 0 in supplemented RIPA after treatment with freshly prepared sodium nitroprusside (100 nM) for 5 min to induce nitric oxide (NO)-mediated signaling. (D) Protein from VSMC was probed for (E) vasodilator-stimulated phosphoprotein (VASP) and (F) phospholamban (PLN) phosphorylation. Statistical comparisons were made with the Student’s t-test between WKY and SHR or SHR and SHR + PBD (n = 6/group). Data are expressed as mean ± SD.

### A plant-based diet differentially impacted redox status and inflammation

Oxidative stress is a key factor in driving EC and VSMC dysfunction. This led us to explore whether proteins involved in the redox pathway were impacted. Superoxide-neutralizing proteins, superoxide dismutase (SOD)1 and SOD2, were significantly increased in LV tissue in SHRs consuming PBD compared to SHRs consuming the control diet (Figure 11A–C). Similarly, catalase (CAT), a hydrogen peroxide-neutralizing protein, was also significantly increased in SHRs consuming PBD, but glutathione peroxidase (GPx1) was not (Figure 11A, D, E). Nuclear factor erythroid 2-related factor (NRF)2, a master regulator of endogenous antioxidant defense, was significantly reduced in nuclear LV extracts in SHRs consuming the control diet compared to WKY, and SHRs consuming PBD (Figure 12A, B). Nuclear extraction purity was validated (Figure S8) with the expression of nuclear marker, specific protein (SP)1, in nuclear and cytosolic extracts. Glutathione is known to be regulated by NRF2, and reduced glutathione was also significantly increased in LV tissue of SHRs consuming PBD compared to SHRs consuming the control diet (Figure 11C). In isolated cells (54.3% VSMC, 45% fibroblast), the pro-oxidant-associated NADPH-oxidase subunit, p22phox, was significantly reduced in SHRs consuming PBD versus SHRs consuming the control diet, as was the lipid peroxidation marker, 4-hydroxy-2-nonenal (4-HNE) (Figure S9). These results suggest that PBD improved cardiac oxidative stress management.

**Figure 11.**
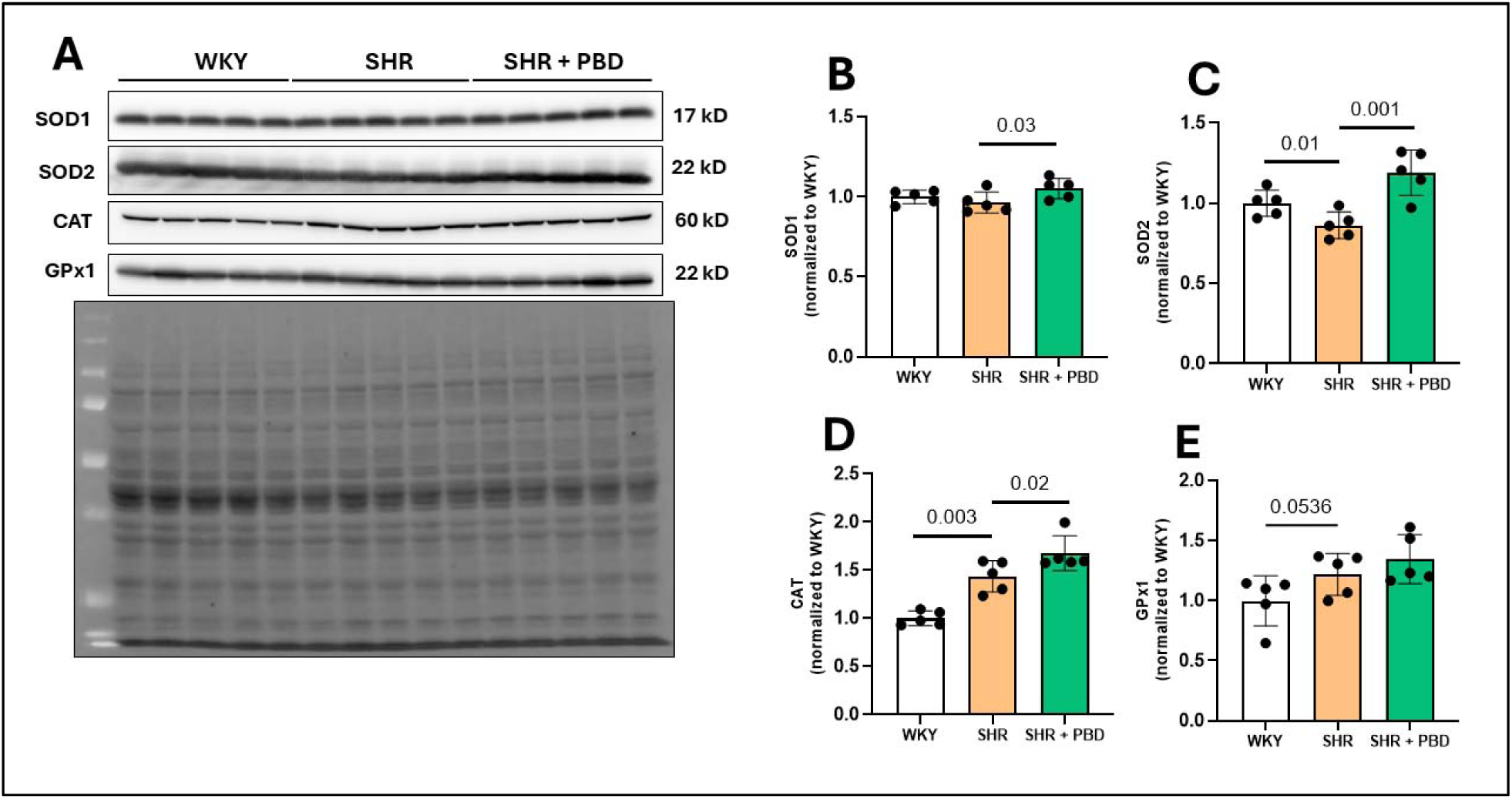
Cardiac endogenous antioxidant defense. Wistar-Kyoto (WKY) rats and spontaneously hypertensive rats (SHR) consumed a purified diet for 24 weeks. After 24 weeks, a subset of SHRs were switched from the control diet to PBD, while a subset of SHRs and WKY continued the control diet for 12 additional weeks. (A) Left ventricular protein was isolated and (B) superoxide dismutase (SOD)1, (C) SOD2, (D) catalase (CAT), and (E) glutathione peroxidase (GPx)1 was probed via western blot. Statistical comparisons were made with Student’s t-test between WKY and SHR or SHR and SHR + PBD (n = 5/group). Data are expressed as mean ± SD.

**Figure 12.**
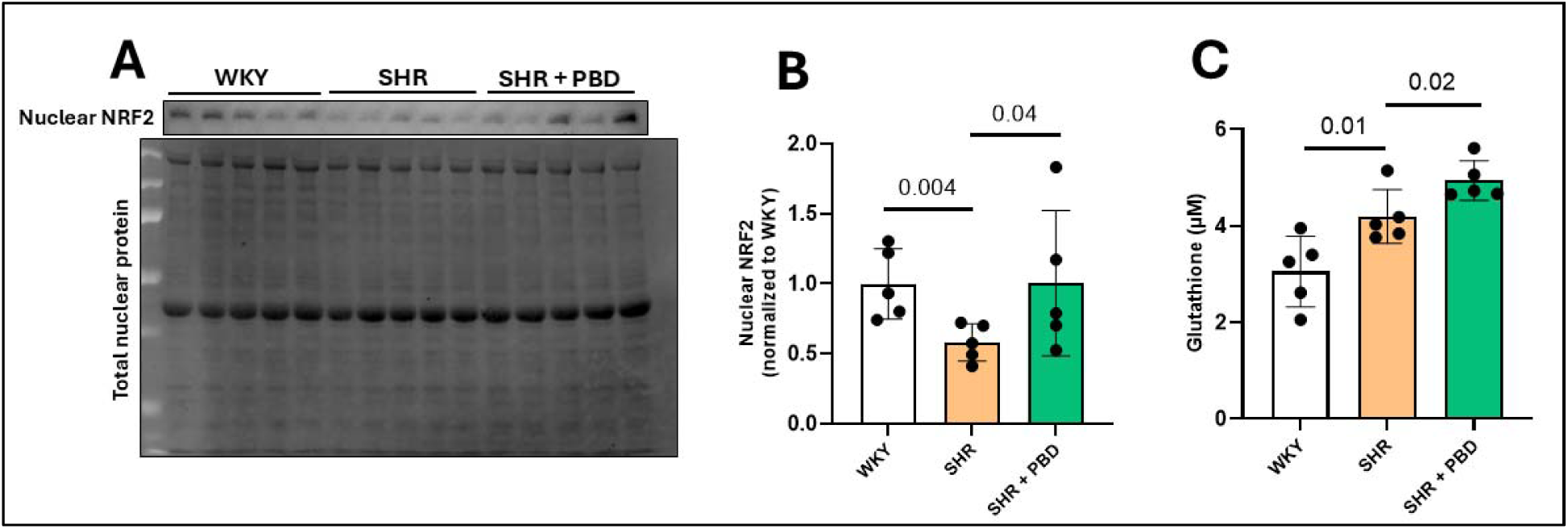
Cardiac antioxidant regulation. Wistar-Kyoto (WKY) rats and spontaneously hypertensive rats (SHR) consumed a purified diet for 24 weeks. After 24 weeks, a subset of SHRs were switched from the control diet to PBD, while a subset of SHRs and WKY continued the control diet for 12 additional weeks. (A) Left ventricular nuclear protein was isolated and (B) nuclear factor erythroid 2-related factor 2 (NRF2) was probed via western blot. (C) Glutathione concentrations in the LV were determined with a commercially available kit. Statistical comparisons were made with Student’s t-test between WKY and SHR or SHR and SHR + PBD (n = 5/group). Data are expressed as mean ± SD.

Next, we examined the impact of PBD on cardiac inflammatory signaling, anticipating that PBD might reverse detrimental changes and/or enhance beneficial counter-regulatory changes that had ensued in SHRs. Expression of phospho-p65, which is known to reflect activation of canonical NF-κB signaling, followed the latter pattern in that it was lower in SHRs, relative to WKYs, and further reduced in PBD-fed SHRs (Figure 13A, B). In contrast, levels of phospho-p38 were lower in SHRs compared to WKYs but were restored by PBD (Figure 13A, C) and SHRs consuming PBD. Analogous to results for phospho-p65, expression of phospho-SAPK/JNK and c-Jun were lower in SHRs and even lower in PBD-fed SHRs although only the differences between SHRs fed the control and PBD reached statistical significance (Figure 13A, C, D). While p38 and SAPK/JNK both can regulate c-Jun^29, 30^, SAPK/JNK appeared to be a more specific regulator in this context. PBD did not attenuate pro-oxidative and inflammatory processes in isolated cardiac white blood cells (Figure S10), nor did we observe significant effects of PBD in the hypothalamus, a region of the brain involved in mediating sympathetic output in hypertension^31^ (Figure S11). These results are in accordance with reduced cardiac inflammatory signaling playing a role in mediating benefits of a PBD.

**Figure 13.**
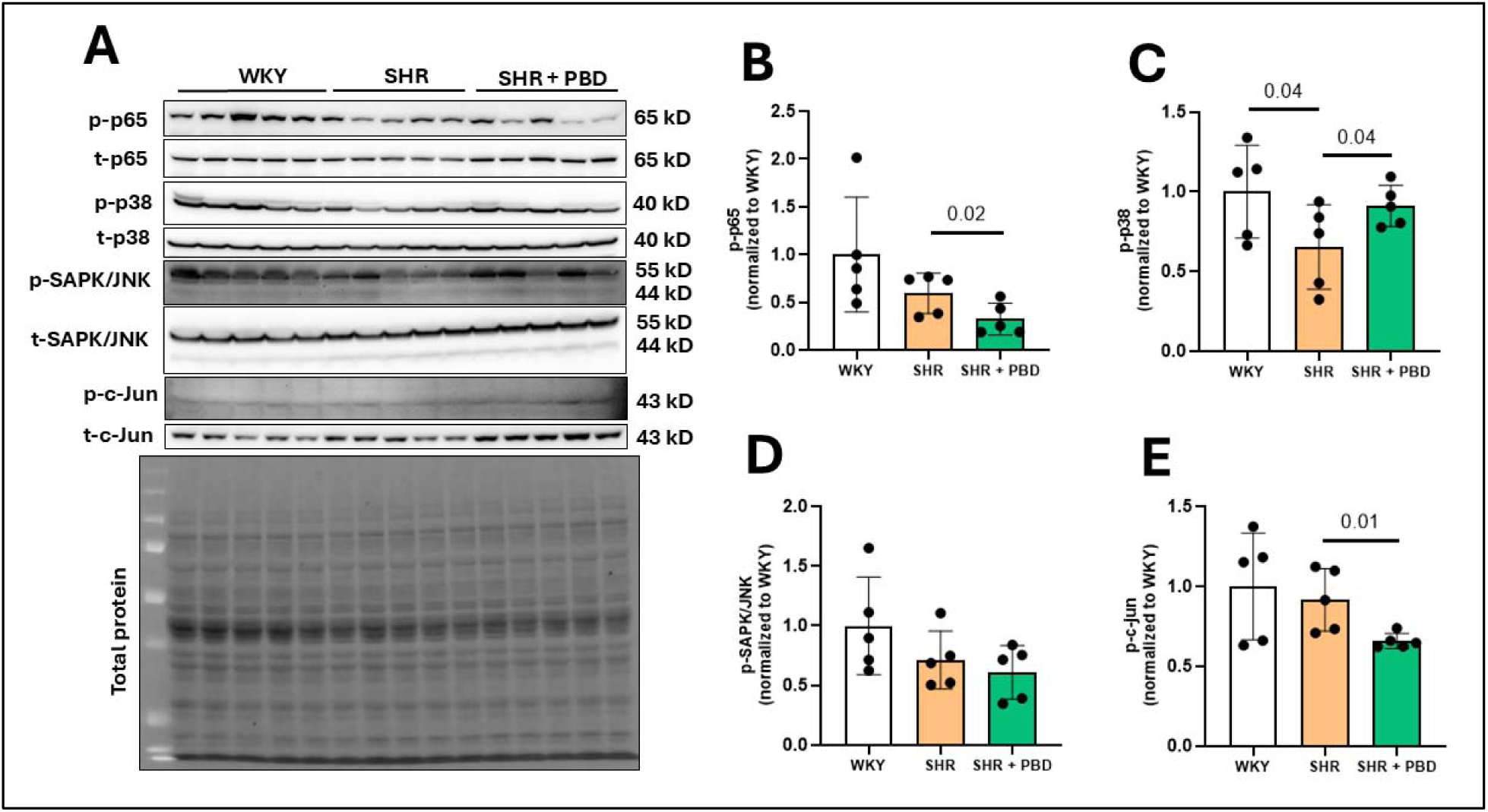
Cardiac inflammatory signaling. Wistar-Kyoto (WKY) rats and spontaneously hypertensive rats (SHR) consumed a purified diet for 24 weeks. After 24 weeks, a subset of SHRs were switched from the control diet to PBD, while a subset of SHRs and WKY continued the control diet for 12 additional weeks. (A) Left ventricular protein was isolated and probed for (B) p38 mitogen-activated protein kinase (MAPK), (C) stress-activated protein kinases (SAPK)/Jun amino-terminal kinases (JNK), (D) c-Jun, and (E) p65, a nuclear factor-kappa B (NF-κB) subunit phosphorylation. Statistical comparisons were made with Student’s t-test between WKY and SHR or SHR and SHR + PBD (n = 5/group). Data are expressed as mean ± SD.

### A plant-based diet treated LV fibrosis in CMD

Fibrosis is a cause, consequence, and amplifier of hypertension-induced cardiac dysfunction. While analysis of cardiac tissue via hematoxylin and eosin staining did not reveal abnormalities in SHRs (Figure S12), this did not rule out the possibility of subtle, early-stage fibrosis. Accordingly, we investigated this notion by visualizing collagen via Sirius red staining of coronal sections of the LV. Despite not having reached definitive levels of heart failure, collagen-dense regions were apparent in SHRs but not in WKYs at 24 weeks of age. Accordingly, quantitation of collagen staining indicated over a 100% increase in Sirius red staining in SHRs relative to WKYs. Meanwhile, PBD prevented fibrosis, nearly completely, by these measures. Furthermore, a very similar pattern of results was observed at 40 weeks even though the PBD was only initiated at 24-weeks of age, indicating that PBD not only prevented, but also reversed early-stage fibrosis in SHRs.

**Figure 13.**
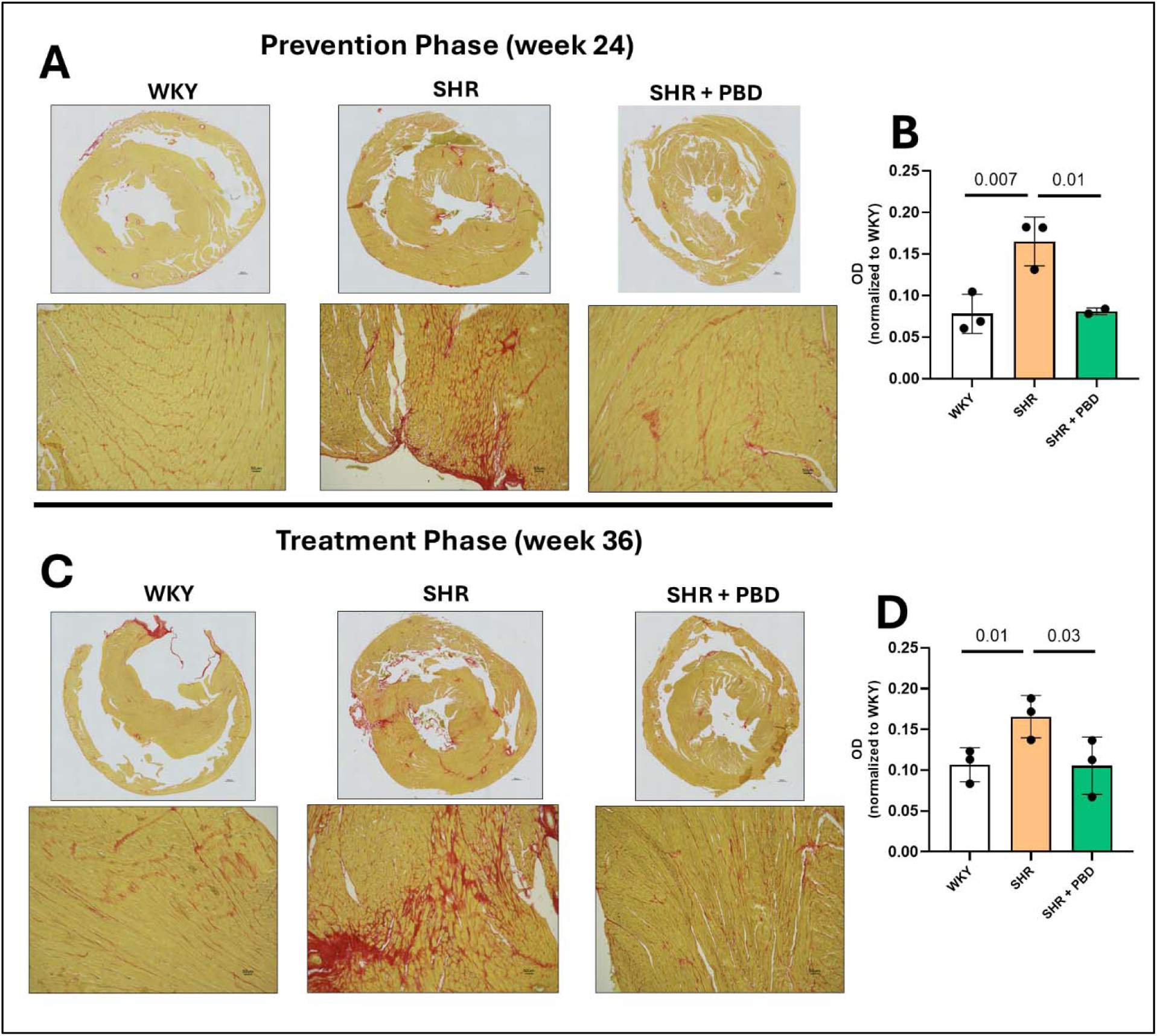
Left ventricular fibrosis. Wistar-Kyoto (WKY) rats or spontaneously hypertensive rats (SHR) consumed a purified diet or 28% plant-based diet (PBD) for 24 weeks with or without antibiotics (ABX). After 24 weeks, a subset of SHRs were switched from the control diet to PBD, while a subset of SHRs and WKY continued the control diet for 12 additional weeks. Hearts were processed for histological analysis and stained for collagen with Sirius red dye. Areas of peak fibrosis (10x magnification) were used for quantification. Blood vessels and collagen attachment sites were avoided for the analysis. Data were analyzed with ImageJ and FIJI plugin. Statistical comparisons were made with Student’s t-test between WKY and SHR or SHR and SHR + PBD (n = 2-3/group). Data are expressed as mean ± SD.

## 4. Discussion

Hypertension is a predominant risk factor and an important contributor to CMD. Accordingly, we hypothesized that SHRs might develop CMD and thus serve as a clinically relevant model of this disorder. We report confirmation of this hypothesis. Furthermore, our use of this tractable model to investigate the impacts of diet on CMD revealed that a PBD both prevented and reversed CMD. Such benefits of a PBD occurred without alleviation of hypertension, a finding we did not anticipate, but which may have reflected that the congenital hypertension of SHRs is difficult to alleviate. In any case, it highlights the potential of PBDs to mitigate CMD irrespective of impacts on blood pressure. The beneficial effects of the PBD correlated with, and were likely mediated by, both endothelial- and VSMC-dependent mechanisms (Figure 14).

**Figure 14.**
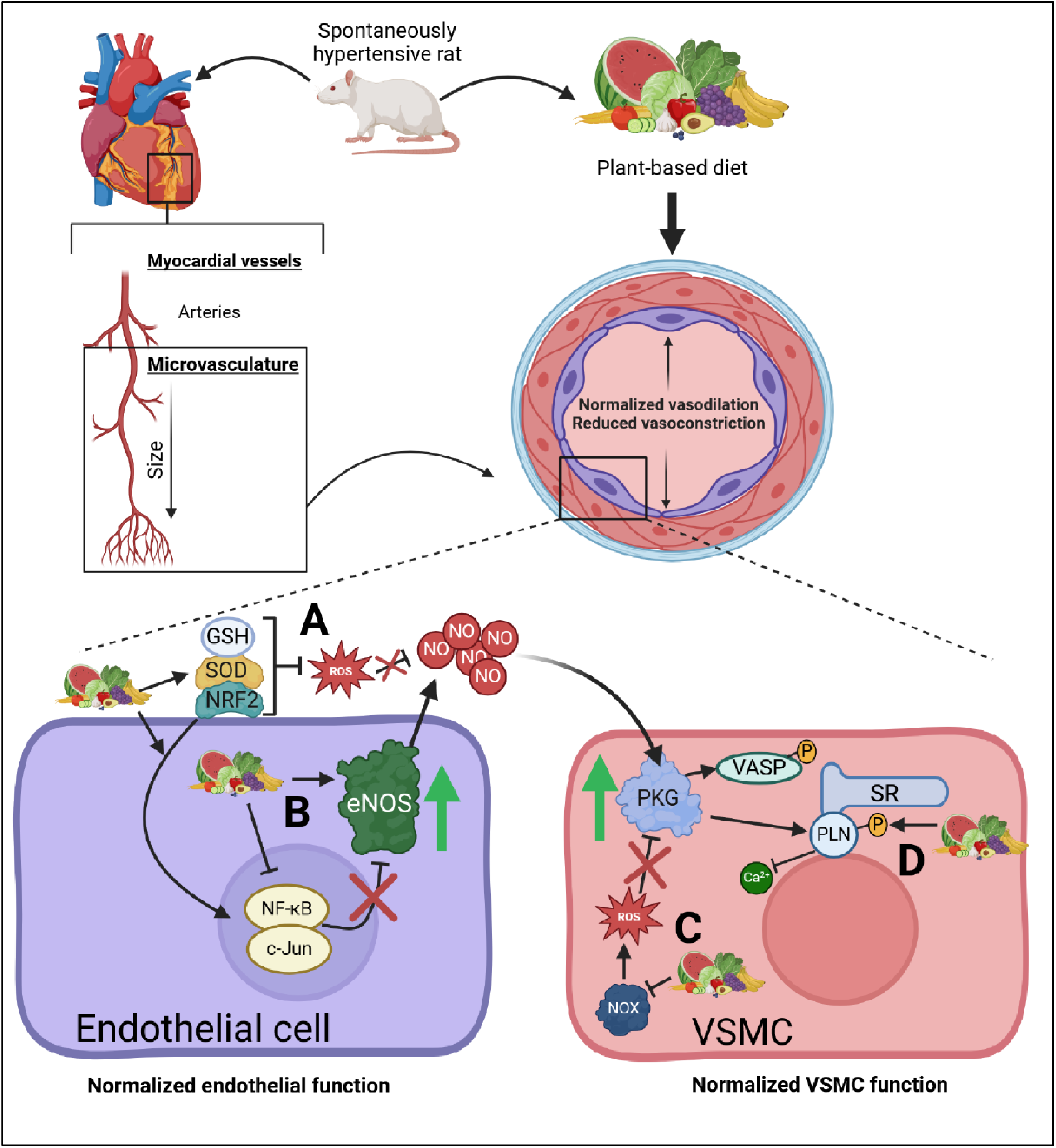
Model by which a PBD targets coronary microvascular dysfunction (CMD) in hypertension. Notable cellular differences within the microvasculature of the cardiac ventricles of spontaneously hypertensive rats (SHR)s fed a plant-based diet (PBD) compared to a control diet are illustrated. (A) PBD enhanced endogenous antioxidant defense by increasing superoxide dismutase (SOD), glutathione (GSH), and increasing nuclear translocation of nuclear factor erythroid 2-related factor (NRF)2. The upregulation of these proteins leads to the direct and indirect neutralization of reactive oxygen species (ROS), preserving nitric oxide (NO) bioavailability. (B) a PBD also enhanced and preserved endothelial nitric oxide synthase (eNOS) expression and function in the endothelium, a protein which synthesizes NO. A PBD also reduced the expression of pro-inflammatory transcription factors, nuclear factor kappa B (NF-κB) and c-Jun, which can downstream reduce eNOS expression and activity. (C) protein kinase G (PKG) activity was notably increased in VSMCs due to increased vasodilator-stimulated phosphoprotein (VASP) phosphorylation. Activation of PKG can be impeded by increased ROS, and a PBD reduced the expression of p22phox, a subunit of NADPH-oxidase (NOX) enzymes which produce ROS, potentially (D) as a consequence of improved PKG activity, a PBD was able to increase phospholamban (PLN) phosphorylation of the sarcoplasmic reticulum (SR), a protein involved in sequestration of calcium ions, resulting in vascular smooth muscle cell (VSMC) relaxation.

VSMC relaxation is driven by NO, which is produced by eNOS. Thus, increased VSMC relaxation in PBD-fed animals may have reflected increased NO and/or increased responsiveness of VSMC to NO. A PBD resulted in increased eNOS phosphorylation (Figure 10A, C), indicating increased eNOS activity. Indeed, the L-NAME-induced reduction in myocardial blood flow, which reflects loss of eNOS activity, was reduced in SHRs, compared to WKYs, and restored by PBD-feeding, suggesting increased eNOS activity in PBD-fed SHRs (Figure 9). Additionally, a PBD resulted in increased levels of serum NO metabolites (Figure S3). While endogenous production of NO via eNOS likely contributed to this quantity, it cannot fully explain these findings, since WKY had preserved eNOS and did not have significantly increased NO metabolites. Thus, changes in NO metabolites likely reflect that the PBD is rich in nitrates. This may have also contributed to increased NO generation via eNOS^32^. Collectively, these data indicate impaired endothelial function which was rescued by PBD consumption. Consistent with this mechanism, reversal of CMD by PBD via 12-week course of PBD feeding restored NO-mediated signaling in VSMCs, which concomitantly led to increased PLN phosphorylation (Figure 10D, F). Thus, both the endothelium and VSMCs were favorably impacted by the PBD. These findings may have high translatability to clinical settings wherein CMD needs to be treated rather than merely be prevented.

Preservation of endothelial and VSMC function may have been a consequence of an improved redox state, as endogenous antioxidant defense was upregulated by PBD (Figure 11-12; Figure S9). Increased oxidative stress and inflammation can facilitate eNOS uncoupling, reducing NO availability, resulting in endothelial dysfunction. Similarly, PKG signaling as a result of NO can be impaired in the presence of oxidative stress and inflammation, resulting in increased VSMC contraction^8^. Oxidative stress and inflammation can also drive pro-fibrotic processes^33^, which was attenuated and reversed in the LV by the consumption of a PBD in the present study (Figure 13). While fibrosis is not classically thought of as reversible, early stage fibrosis may be^34^, as was the case in the present study. Indeed, extensive LV fibrosis would be detrimental to cardiac function and lead to heart failure. Heart failure does not typically manifest in female SHRs until 24 months of age^27^, thus, an extension of this study may have resulted in more pronounced fibrosis.

The persistence of hypertension in PBD-fed SHRs was likely due to reduced eNOS in the conduit blood vessels as represented by aortic and mesenteric protein (Figure S6); however, this was not the case in the coronary microvascular blood vessels. Indeed, we believe that there may be distinct effects between conduit blood vessels and organ microvasculature. We showed that in liver and kidney, eNOS is diminished in SHRs, which is attenuated by a PBD (Figure S7), indicating that the microvasculature was preserved in the organs. Conduit blood vessels are involved in regulating blood pressure due to their elastic nature, and these appeared to be compromised based on eNOS protein data unlike the microvasculature. It is also important to consider that there are multifaceted mechanisms within the SHR that drive hypertension. These include structural abnormalities of the vasculature, VSMC hyperreactivity, overactive sympathetic output, as well as renin-angiotensin system upregulation ^35^. These factors may make dietary changes insufficient in overcoming this pronounced hypertensive phenotype. In contrast in humans, we have shown that a PBD could treat hypertension in a mostly female cohort of patients and significantly reduce the need for blood pressure-managing medications^36^. Thus, it is noteworthy that in the present model, CMD was attenuated and reversed despite hypertension.

Considering the essential role of the gut microbiota in metabolizing phytochemicals, such as polyphenols, it was expected that any beneficial effects of the PBD would be negated by ABX. In contrast, a PBD improved CMD and endothelial function even with ABX (Figures 7 & 9). Parent phytochemical compounds are in much lower concentrations in serum following the consumption of phytochemical-rich foods compared to their metabolites. These compounds undergo various hydrolysis, deglycosylation, dehydroxylation, and demethylation reactions facilitated by gut microbiota to increase absorption^13^. Indeed, unique polyphenol profiles in serum can be observed 2 h following the consumption of red raspberry compared with 24 h which closely tied with improved macrovascular function at both time-points^37^. The consumption of a single 500 mg dose of cyanidin-3-glucoside, a common anthocyanin found in blueberries and blackberries, resulted in the presence of 33 different metabolites, each with distinct times of appearance in serum at 1, 6, and 24 h post-consumption^38^. These postprandial profiles independently were effective at physiological concentrations in attenuating endothelial cell inflammation^39^. Further, gut microbe populations have differing metabolic capabilities, with *Bacteroides* conferring negative health effects compared with *Prevotella* which are beneficial to health^40^. Indeed, this favorable shift occurs rapidly within 24 h of consuming a PBD in humans, and reverses once the PBD is ceased^41^. Thus, polyphenols are known for modulating microbial populations^13^. Microbial analysis in the present study revealed increased *Prevotella* in SHRs consuming a PBD at week 24 (Figure 2C), a microbe associated with PBDs^42^. Nonetheless, we cannot draw firm conclusions regarding the role the microbes in mediating benefits of PBD in that ABX treatment did not lead to clear differences in CMD. However, it should be noted that depletion of microbes in the SHR may lead to some benefit, albeit we were not powered to detect this difference statistically. Mean CFR values for SHR and SHR + ABX were 1.7 and 2.0, respectively. Other studies have identified that harmful microbes may exist in SHRs. For example, fecal transplantation from SHRs to WKY (all males), and vice versa, results in an increase in blood pressure in WKY and a reduction of blood pressure in SHRs ^43^. Thus, future studies should assess whether harmful microbes may also be impacting coronary microvascular circulation.

Several limitations in the present investigation exist. Firstly, it is known in rodents, such as mice, that isoflurane can induce a hyperemic response^44, 45^. However, other investigators have used isoflurane induction in rats followed by injection with adenosine or dobutamine, followed by CFR measurement with echocardiography or PET scan^46, 47^. We performed preliminary studies and found that 1% isoflurane was unable to maintain the anesthetic plane after induction, and coronary velocity was artificially elevated since animals were becoming alert. At 2.5% isoflurane, coronary velocity dropped significantly, and the anesthetic plane was maintained. Thus, we used 2.5% isoflurane for our CFR measurements, and all animals were maintained at a similar anesthetic depth with the same isoflurane concentration. Secondly, while we aimed for this research to be as clinically relevant as possible, it is unclear whether the unique genetic phenotype of the SHR is translatable to humans. For example, angiotensin II infusion and Dahl salt sensitive rats may result in different outcomes. Furthermore, this animal model of CMD is newly established^23^, thus, more established models of CMD utilizing metabolic syndrome characteristics^48^ require investigation. Another limitation is the lack of purity in the VSMC culture. Initially, one aim of this study was to isolate ECs. However, these cells, despite being CD31^+^ and CD90^−^, would not successfully culture. It was identified that VSMCs could be isolated with these methods, however. Initial labelling with MYH11 antibody (50-6400-80, ThermoFisher) a VSMC-specific marker, with flow cytometry revealed ∼99% purity. However, it was later discovered that this antibody was nonspecific and of poor quality. Additional testing with α-SMA and CD90 antibodies revealed that the VSMC culture was contaminated with fibroblasts (Figure S5). Since this VSMC data had already been collected, it was not possible to go back and remove fibroblasts with CD90 magnetic beads. Nonetheless, since PLN is VSMC-specific, we are confident that we have captured the relevant data illustrating preserved VSMC function with SNP. However, VASP phosphorylation can occur in fibroblasts as well, and thus, is a limitation in our interpretations of VSMC data. An additional limitation is that we did not assess gut microbial profiles of ABX animals. However, since bacteria in feces were reduced 10,000-fold in ABX animals and considering that ABX treatment did not diminish CFR in PBD-supplemented animals, we did not find it relevant to perform 16s rRNA sequencing on ABX feces. Nonetheless, it is possible that these microbes in ABX, while minute, could still have some physiological role. Thus, future studies should assess these parameters. As emphasized throughout, women are more likely to be impacted by CMD, particularly those that are post-menopausal. However, since we used cycling female rats, this does not fully reflect the human phenotype. Further, since we only used female animals in this study, sex-differences could not be investigated. We focused this study on female rats and therefore could not investigate sex-differences. While CMD occurs in both men and women, and is prognostic in both sexes, women are more impacted with CMD with more angina and lower quality of life. In the setting of non-obstructive CAD, Women have more serve CMD compared to men. Thus, this phenotype reflects the SHR more closely, since SHRs do not develop atherosclerosis spontaneously, and are even resistant to diet-mediated hypercholesterolemia-induced atherosclerosis^49^. Future studies should evaluate the effects of PBD in both sexes, especially those in older animals.

In conclusion, a PBD was able to both prevent CMD development and reverse CMD once established in an animal model of hypertension. These effects were likely due to improved endothelial function and possibly improved VSMC function. This occurred independent of hypertension. Pilot clinical studies evaluating the effects of a PBD in CMD are warranted.

## Data availability statement

Raw western blot images are provided on Figshare, an online repository with the following DOI: 10.6084/m9.figshare.28790810. Microbiota 16s sequencing data can be found at the following DOI on Figshare: 10.6084/m9.figshare.28781054. Other data can be provided by the corresponding author upon reasonable request.

## Ethical Approval

This study was approved by the Institutional Animal Care & Use Committee (IACUC) at Georgia State University. The approval number for this study is A23025.

## CRediT authorship contribution statement

**Rami S. Najjar**: Writing – review & editing, Writing – original draft, Visualization, Methodology, Investigation, Formal analysis, Conceptualization, Funding acquisition, Project administration. **Khan Hekmatyar**: Writing – review & editing, Methodology, Investigation. **Yanling Wang**: Writing – review & editing, Methodology, Formal analysis. **Vu Ngo**: Writing – review & editing, Methodology, Formal analysis. **Hannah L. Lail**: Writing – review & editing, Methodology. **Juan P. Tejado**: Writing – review & editing, Methodology, Formal analysis. **Jessica P. Danh**: Writing – review & editing, Investigation. **Desiree Wanders**: Writing – review & editing, Resources. **Rafaela G. Feresin**: Writing – review & editing, Resources. **Puja K. Mehta**: Writing – review & editing, Methodology. Conceptualization. **Andrew T. Gewirtz**: Writing – review & editing, Funding acquisition, Methodology, Supervision, Resources.

