## Supplementary Figure for "Prevention and reversal of hypertension-induced coronary microvascular dysfunction by a plant-based diet"

**Table S1.** Composition of the modified control diet (D23061303) and plant-based diet (D23061204).

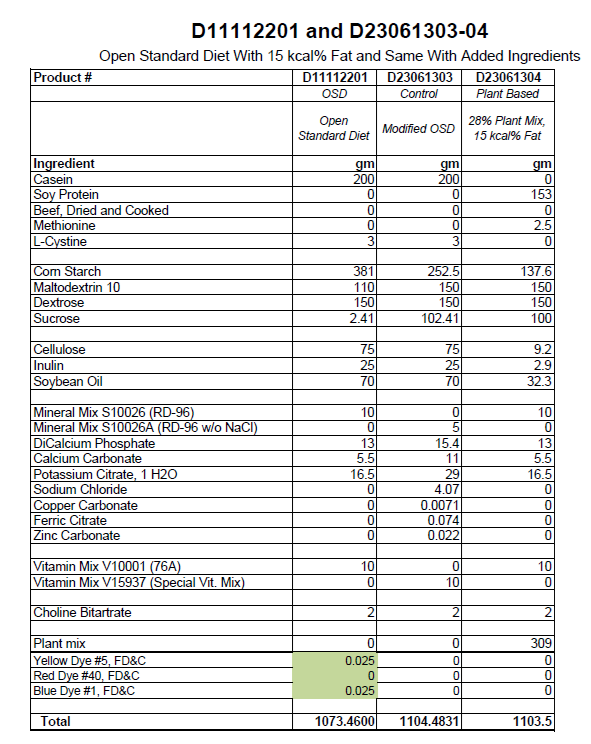

**Table S2.** Macronutrient breakdown of the modified control diet (D23061303) and plant-based diet (D23061204).

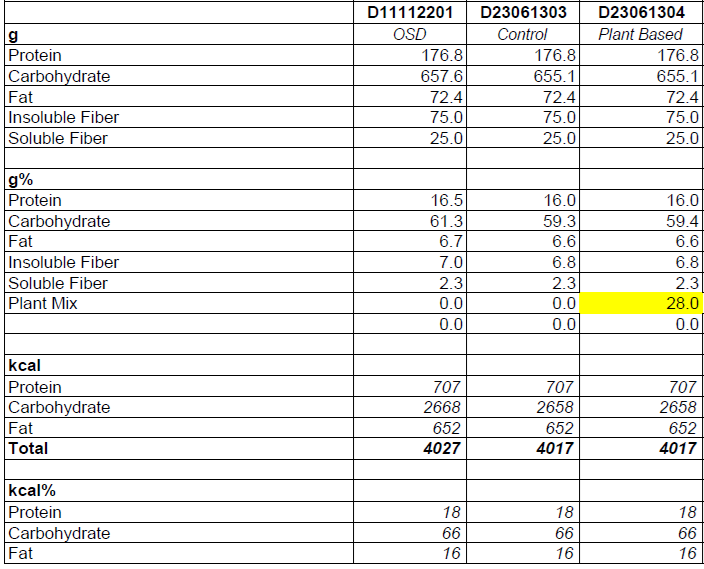

**Table S3.** Body weight at weeks 28 and 40 weeks of age. Average body weight of 2 rats/cage (n = 8-14). Data are presented as mean ± standard deviation. Data presented are truncated to the nearest whole number.

| **Age of animals** | **WKY** | **SHR** | **SHR + PBD** | **SHR + ABX** | **SHR + ABX + PBD** |
| --- | --- | --- | --- | --- | --- |
| **Week 28** | 238 ± 10 g | 207 ± 5 g | 201 ± 7 g | 200 ± 2 g | 206 ± 2 g |
| **Week 40** | 244 ± 12 g | 227 ± 8 g | 229 ± 10 g |  |  |

**Table S4.** Systolic (SBP) and diastolic (SBP) blood pressure at weeks 28 and 40 weeks of age (*n* = 5–14/group). Data are presented as mean ± standard deviation. Data presented are truncated to the nearest whole number.

| **Age of animals** | **WKY** | **SHR** | **SHR + PBD** | **SHR + ABX** | **SHR + ABX + PBD** |
| --- | --- | --- | --- | --- | --- |
| **SBP** |  |  |  |  |  |
| **Week 28** | 112 ± 12 | 171 ± 15 | 157 ± 16 | 158 ± 9 | 172 ± 20 |
| **Week 40** | 112 ± 11 | 160 ± 10 | 165 ± 7 |  |  |
| **DBP** |  |  |  |  |  |
| **Week 28** | 71 ± 12 | 122 ± 5 | 105 ± 13 | 106 ± 16 | 107 ± 25 |
| **Week 40** | 80 ± 15 | 107 ± 11 | 111 ± 4 |  |  |

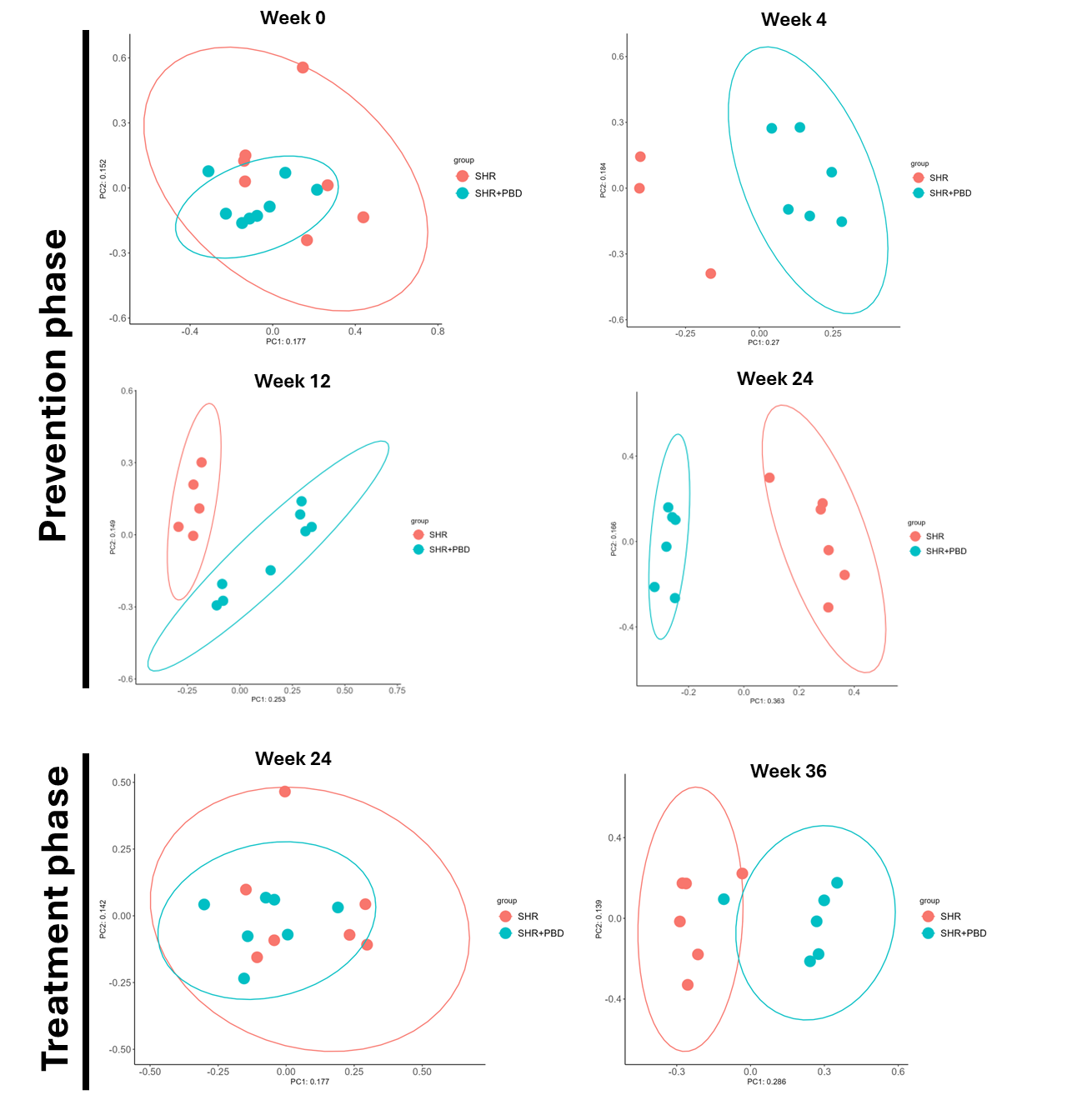

**Figure S1**. **Beta diversity among SHRs consuming a control diet or PBD over time.** Spontaneously hypertensive rats (SHR) consumed a purified diet or 28% (w/w) plant-based diet (PBD) for 24 weeks. After 24 weeks, a subset of SHRs were switched from the control diet to PBD, while a subset of SHRs continued the control diet for 12 additional weeks. PCoA plot of Bray-Curtis distance was shown at different timepoints.

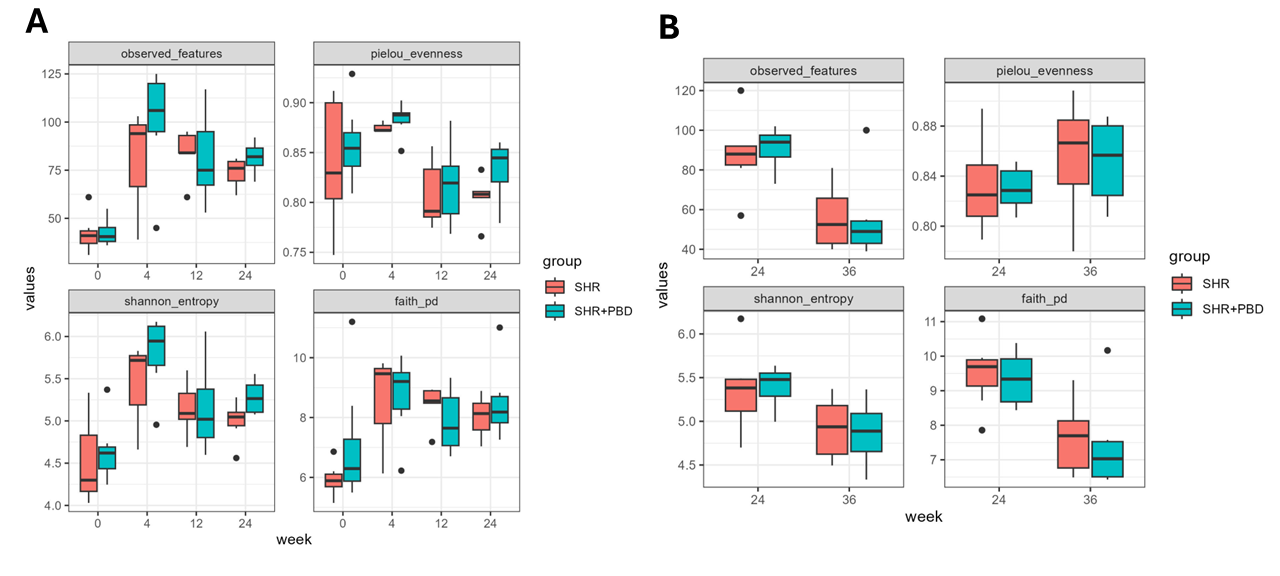

**Figure S2**. **Alpha diversity among SHRs consuming a control diet or PBD over time.** (A) Spontaneously hypertensive rats (SHR) consumed a purified diet or 28% (w/w) plant-based diet (PBD) for 24 weeks. (B) After 24 weeks, a subset of SHRs were switched from the control diet to PBD, while a subset of SHRs continued the control diet for 12 additional weeks. Gut bacteria alpha diversity was measured by four metrics: observed features, evenness, shannon entropy and faith pd.

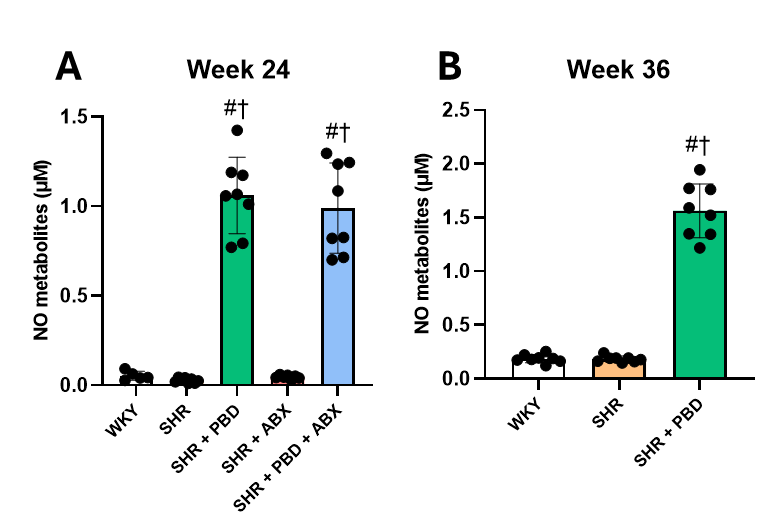

**Figure S3. Serum nitric oxide metabolites (nitrates + nitrites).** Wistar-Kyoto (WKY) rats or spontaneously hypertensive rats (SHR) consumed a purified diet or 28% (w/w) plant-based diet (PBD) for 24 weeks with or without antibiotics (ABX). After 24 weeks, a subset of SHRs were switched from the control diet to PBD, while a subset of SHRs and WKY continued the control diet for 12 additional weeks. Serum was collected and assessed for nitric oxide metabolites at (A) week 24 and (B) week 36. Data was statistically assessed with one-way ANOVA and Dunnett’s post-hoc test (n = 5–8/group). Significance (P ≤ 0.05) is denoted by “#” vs WKY and “†” vs SHR alone. Data are expressed as mean ± SD.

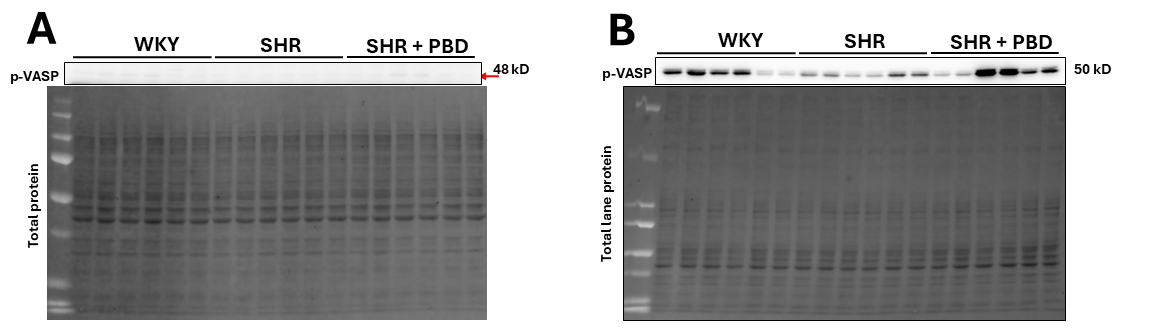

**Figure S4. Validation of sodium nitroprusside function in isolated cells.** Isolated VSMCs were treated with or without sodium nitroprusside (SNP), an agonist of vasodilator-stimulated phosphoprotein (VASP). (A) Expression of phospho-VASP was weak following 300 second chemiluminescent exposure in cells without SNP treatment compared to (B) a much stronger chemiluminescent signal after 5 seconds of exposure in cells treated with SNP. In addition, it was noted that the kD of VASP was slightly less without SNP (48 kD) compared to with SNP (50 kD), which may be a function of post-translational modification increasing the molecular weight.

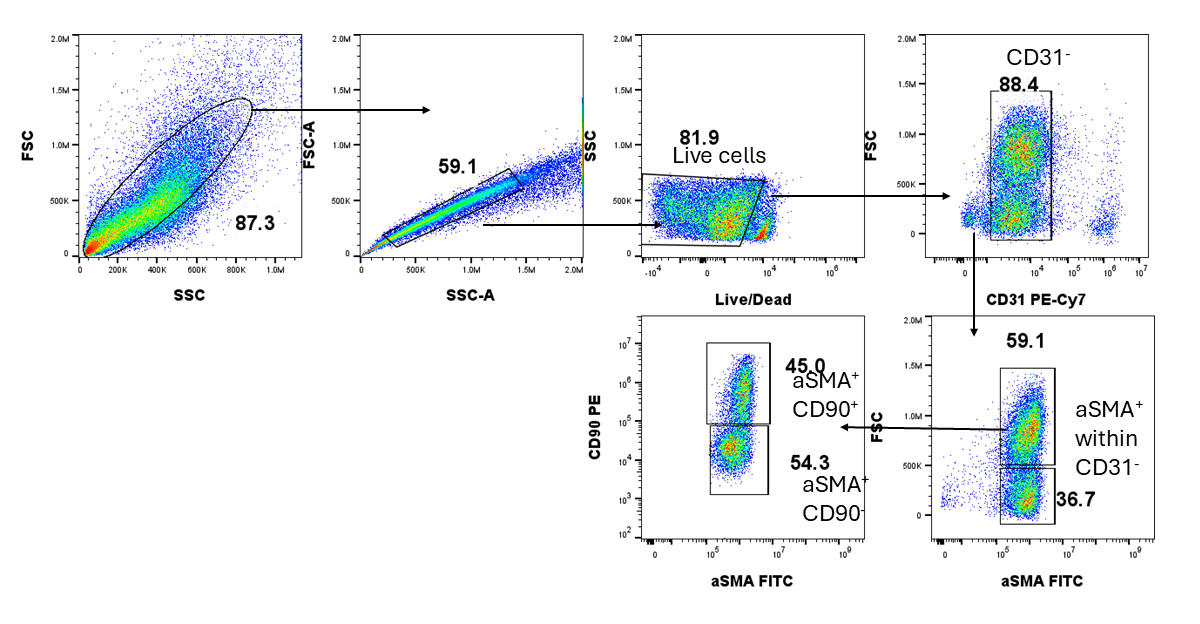

**Figure S5. Gating strategy used to determine cell purity via flow cytometry.** A Wistar-Kyoto (WKY) animal was used in the representative flow cytometry data.

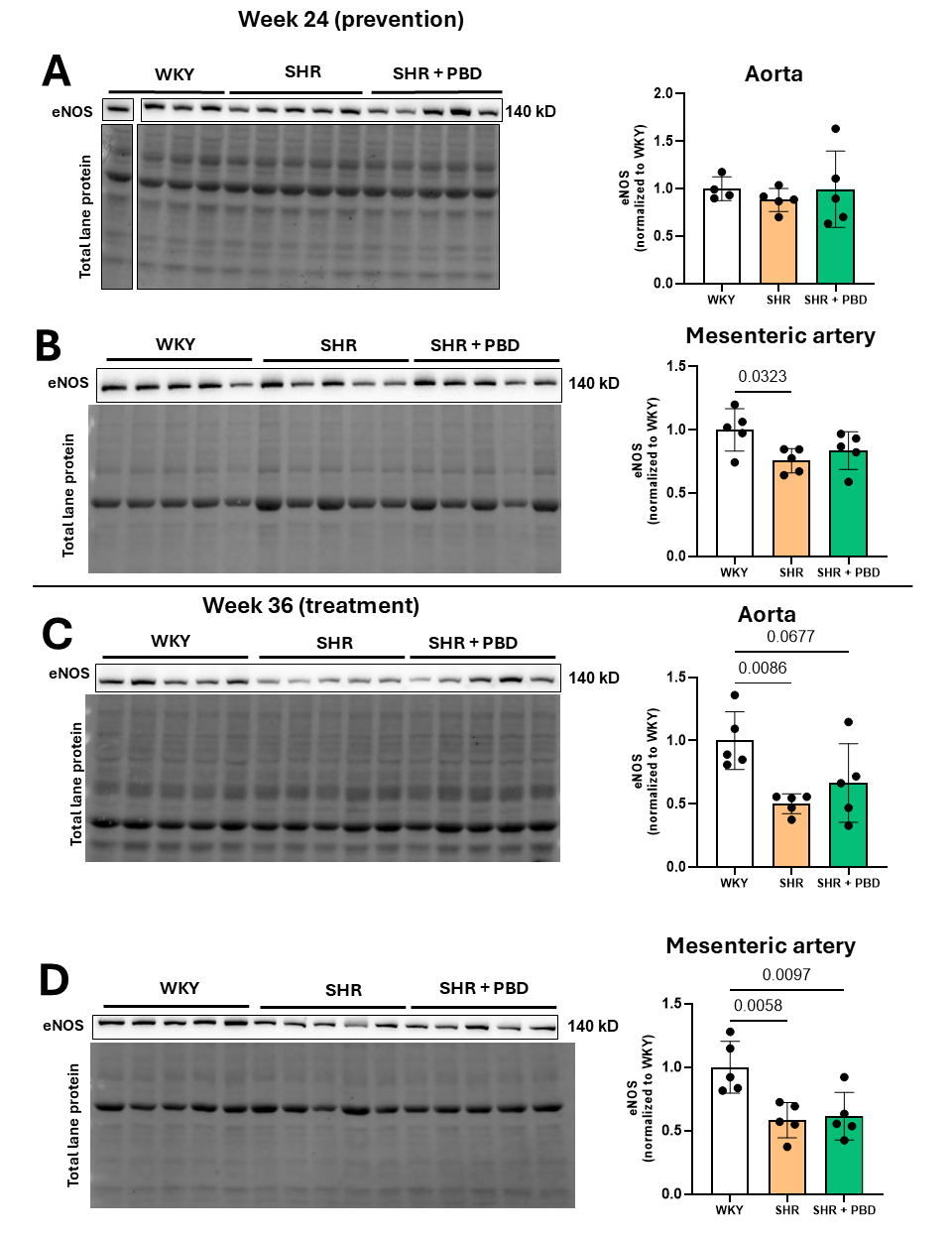

**Figure S6. The protein expression of conduit vessel eNOS in animals at weeks 24 and 36.** Wistar-Kyoto (WKY) rats or spontaneously hypertensive rats (SHR) consumed a purified diet or 28% (w/w) plant-based diet (PBD) for 24 weeks. After 24 weeks, a subset of SHRs were switched from the control diet to PBD, while a subset of SHRs and WKY continued the control diet for 12 additional weeks. Aortas (A, C) and mesenteric arteries (B, D) were excised and assessed for eNOS protein expression. Data was statistically assessed with one-way ANOVA and Dunnett’s post-hoc test (n = 4–5/group).

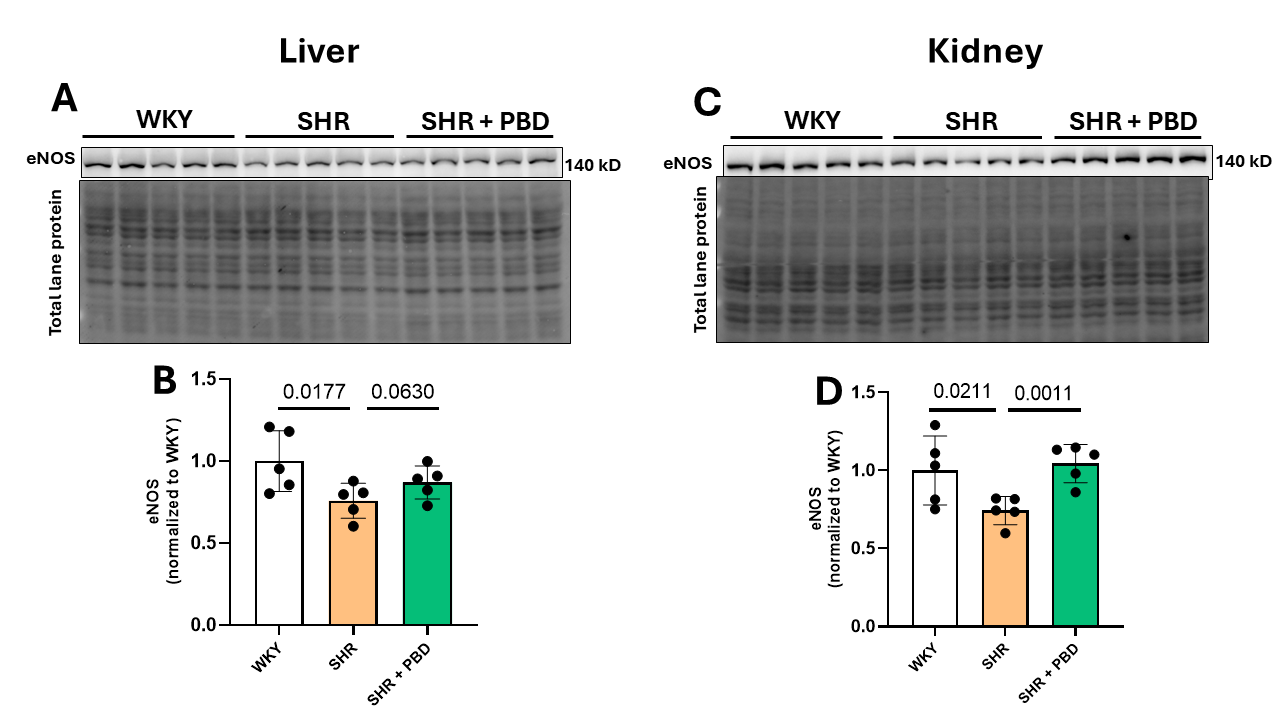

**Figure S7. The protein expression of microvascular organ eNOS in liver and kidney.** Wistar-Kyoto (WKY) rats or spontaneously hypertensive rats (SHR) consumed a purified diet or 28% (w/w) plant-based diet (PBD) for 24 weeks. After 24 weeks, a subset of SHRs were switched from the control diet to PBD, while a subset of SHRs and WKY continued the control diet for 12 additional weeks. Liver (A, C) and left kidneys (B, D) were excised and assessed for eNOS protein expression (n = 4–5/group).

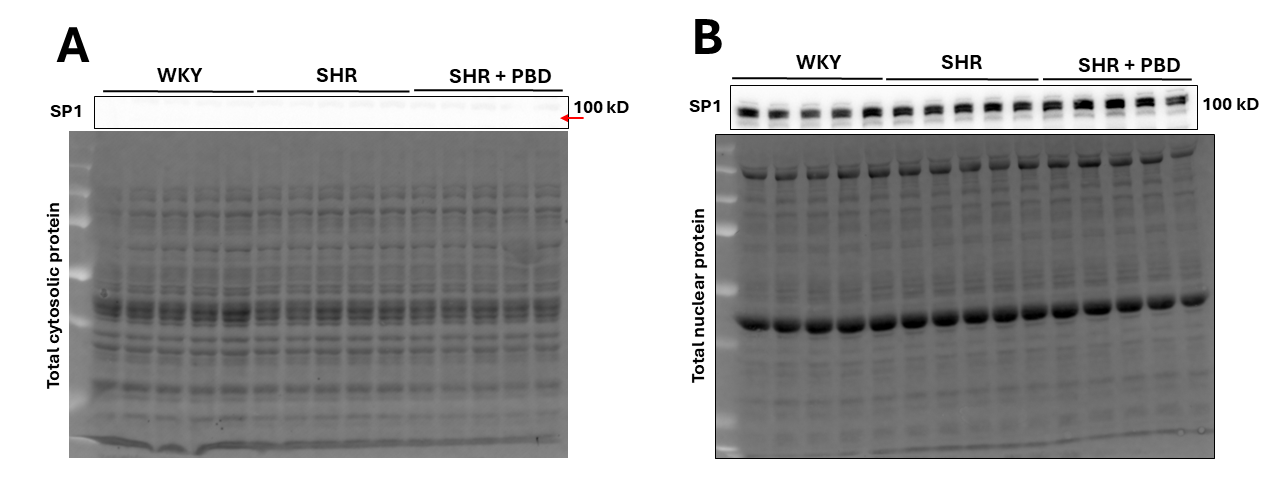

**Figure S8. Validation of nuclear extract purity.** Both (A) cytosolic and (B) nuclear extracts of the left ventricle of the heart were assessed for the nuclear fraction-specific protein, specific protein (SP)1. Following a 2 second chemiluminescent exposure with identical contrast settings, SP1 was absent in the cytosolic extract (A) compared to the nuclear extract (B).

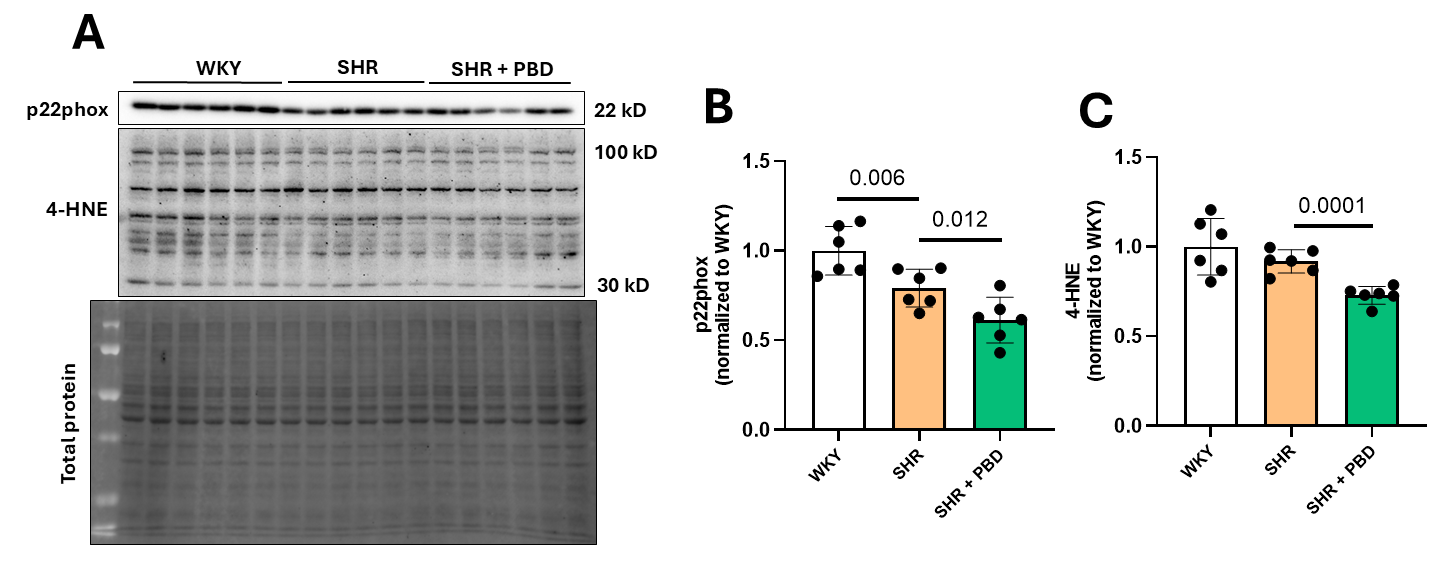

**Figure S9. Oxidative stress markers in isolated VSMCs.** Wistar-Kyoto (WKY) rats and spontaneously hypertensive rats (SHR) consumed a purified diet for 24 weeks. After 24 weeks, a subset of SHRs were switched from the control diet to PBD, while a subset of SHRs and WKY continued the control diet for 12 additional weeks. Vascular smooth muscle cell (VSMC)-comprised cells were isolated from the heart. Following culture and confluence, cells were placed in starvation medium (10% of the growth supplement; 0.5% FBS) for 24 h. Cells were then lysed at passage 0 in supplemented RIPA. (A) Protein from VSMC was probed for (B) p22phox and (C) 4-hydroxy-2-nonenal (4-HNE). Statistical comparisons were made with Student’s t-test (n = 6/group). Data are expressed as mean ± SD.

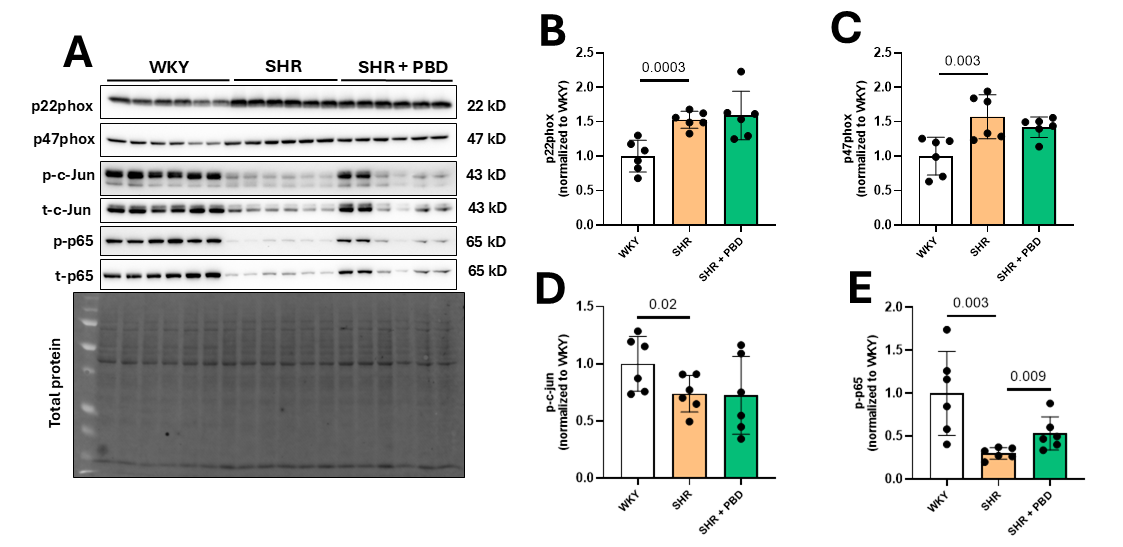

**Figure S10. White blood cell oxidative stress markers and inflammatory signaling proteins.** Wistar-Kyoto (WKY) rats and spontaneously hypertensive rats (SHR) consumed a purified diet for 24 weeks. After 24 weeks, a subset of SHRs were switched from the control diet to PBD, while a subset of SHRs and WKY continued the control diet for 12 additional weeks. White blood cells (WBCs) were isolated from cardiac ventricles using CD45-conjugated magnetic beads. These cells were immediately lysed in supplemented RIPA. (A) Protein from WBCs was probed for (B) p22phox, (C) p47phox, (D) c-Jun, and (E) p65 phosphorylation. Statistical comparisons were made with Student’s t-test (n = 6/group). Data are expressed as mean ± SD.

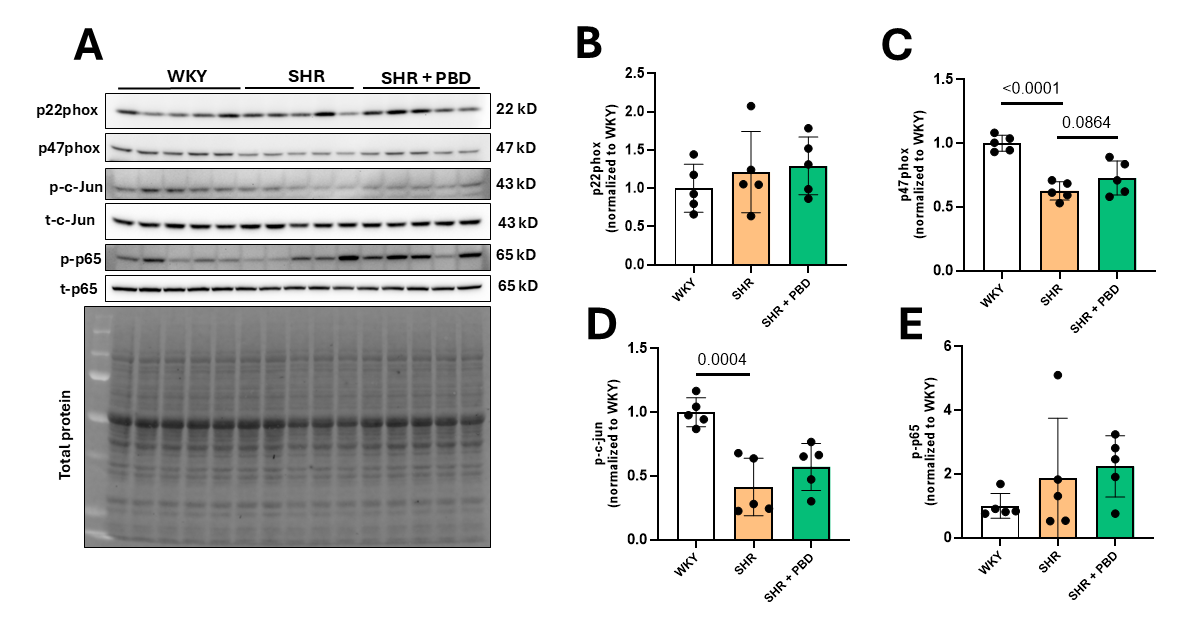

**Figure S11. Hypothalamus oxidative stress markers and inflammatory signaling proteins.** Wistar-Kyoto (WKY) rats and spontaneously hypertensive rats (SHR) consumed a purified diet for 24 weeks. After 24 weeks, a subset of SHRs were switched from the control diet to PBD, while a subset of SHRs and WKY continued the control diet for 12 additional weeks. The hypothalamus was isolated and homogenized in supplemented RIPA. (A) Protein from the hypothalamus was probed for (B) p22phox, (C) p47phox, (D) c-Jun, and (E) p65 phosphorylation. Statistical comparisons were made with Student’s t-test (n = 6/group). Data are expressed as mean ± SD.

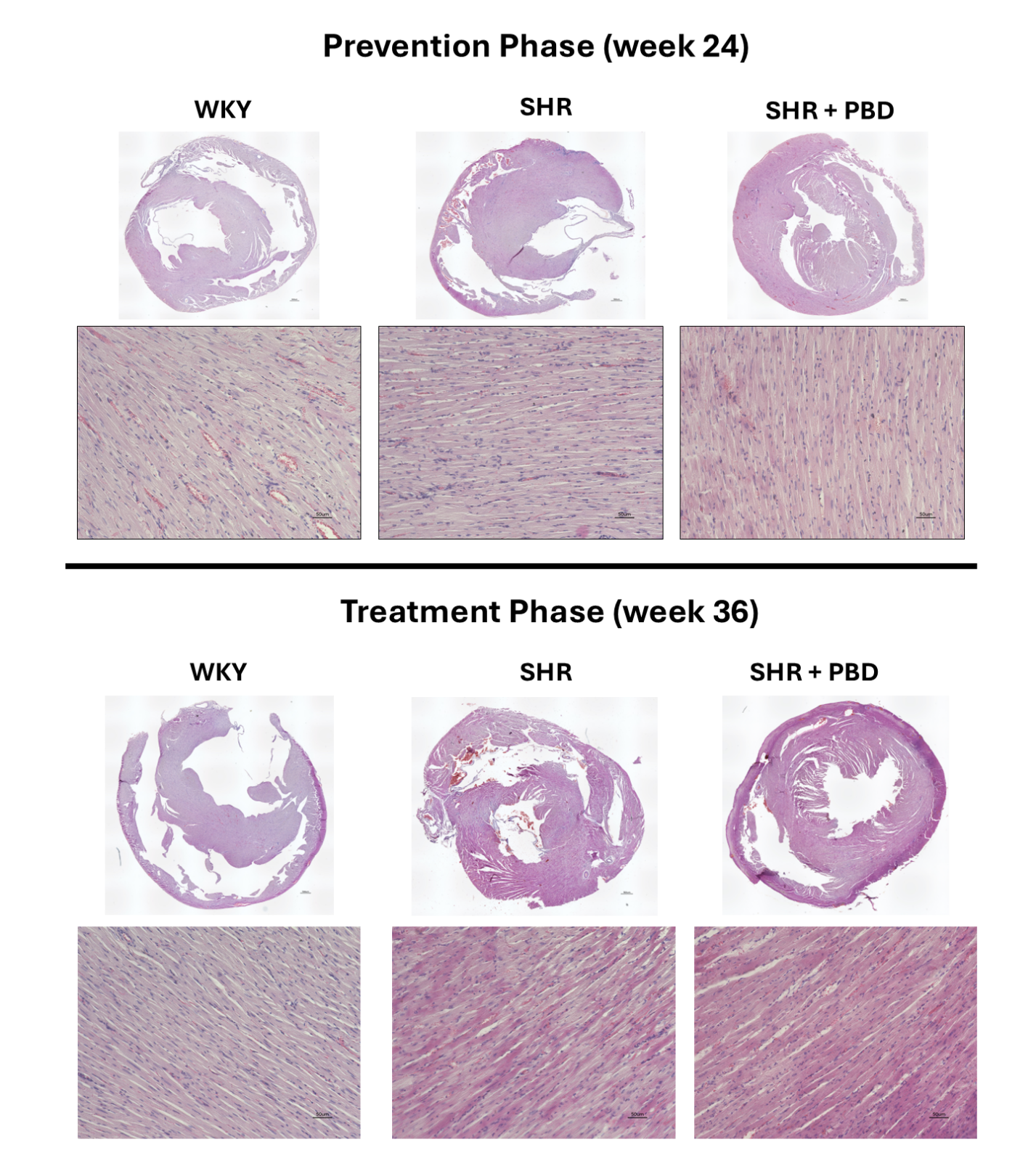

**Figure S12. Histological characteristics of the cardiac left ventricle.** Wistar-Kyoto (WKY) rats or spontaneously hypertensive rats (SHR) consumed a purified diet or 28% (w/w) plant-based diet (PBD) for 24 weeks with or without antibiotics (ABX). After 24 weeks, a subset of SHRs were switched from the control diet to PBD, while a subset of SHRs and WKY continued the control diet for 12 additional weeks. Hearts were processed for histological analysis and stained with H&E. Representative images of animals are shown.
